# *Mycobacterium tuberculosis* manipulates host inflammation and lipid metabolism through the SET1-interacting protein Rv1075c

**DOI:** 10.64898/2026.08.21.746307

**Authors:** Aja K. Coleman, Cory J. Mabry, Morgan J. Chapman, Kaitlyn S. Armijo, Mackenzie H. Smith, Jessica B. Huskey, Lauren Strannahan, Robert O. Watson, Kristin L. Patrick

**Affiliations:** Department of Microbial Pathogenesis and Immunology, Texas A&M University, Naresh K. Vashist College of Medicine, Bryan, TX 77807, USA; Department of Pathology, Microbiology, and Immunology, Division of Molecular Pathogenesis, Vanderbilt University Medical Center, Nashville, TN 37232; Deparentment of Veterinary Medicine and Biomedical Sciences, Texas A&M University, College Station, Texas, 77843; Department of Medicine, Division of Infectious Diseases, Vanderbilt University Medical Center, Nashville, TN 37232

## Abstract

A growing body of literature supports a critical role for nucleomodulins, proteins that traffic to host cell nuclei and manipulate nuclear processes, in intracellular bacterial pathogenesis. Here, we identify the *Mycobacterium tuberculosis* (Mtb) secreted protein Rv1075c as a nucleomodulin that targets a histone modifying protein complex in macrophages. We report that ΔRv1075c Mtb infection elicits a blunted transcriptional response in inflammatory and lipid metabolism pathways and fails to induce foamy macrophage formation in the lungs of infected mice. Using an unbiased mass-spectrometry based approach, we found that Rv1075c interacts with components of the H3K4me3-depositing SET1 histone methyltransferase complex, and that this interaction is required for Rv1075c nuclear localization. Consistent with Rv1075c inhibiting SET1 activity, SET1 deficiency results in hyperinduction of inflammatory genes in activated macrophages. Together, these findings reveal a mechanism by which Mtb engages host chromatin machinery and support a model whereby Rv1075c exploits the SET1 complex to promote a host environment conducive to mycobacterial persistence.

**IMPORTANCE:** Tuberculosis is caused by *Mycobacterium tuberculosis* (Mtb), a remarkably adaptable bacterial pathogen that can survive inside the very immune cells that are supposed to destroy it. To survive, Mtb has evolved ways to interfere with the body’s natural defenses and create conditions that help it persist. Understanding how the bacterium accomplishes this is critical for developing better treatments for TB. In this study, we found that Mtb produces a protein called Rv1075c that travels to the nucleus of infected cells, where it interacts with chromatin remodeling proteins that regulate gene expression. By altering how these genes are controlled, Rv1075c increases the activity of genes involved in inflammation and lipid metabolism, creating a pro-bacterial environment. These findings reveal a previously unknown way that Mtb hijacks host cell gene expression to promote infection, providing new insight into how TB causes disease and suggesting potential targets for future therapies.

## INTRODUCTION

Many bacterial pathogens manipulate host cells by secreting toxins and virulence factors that disrupt immune defenses and promote infection. Increasing evidence suggests that a class of these secreted proteins, known as nucleomodulins, target the host cell nucleus (1, 2). There, nucleomodulins manipulate numerous steps in gene expression (e.g. chromatin remodeling, transcription) and RNA processing (e.g. splicing, mRNA export) (3). The first nucleomodulins were identified in *Agrobacterium tumefaciens*, a soil-dwelling Gram-negative bacterium, that uses VirD2 and VirE4 proteins to traffic bacterial T-DNA into host cells and integrate it into the plant genome (4–6). In recent years, the number of known nucleomodulins targeting mammalian hosts has expanded considerably. Some notable examples include the TAL (transcription activator-like) proteins like AvrBs3, secreted by *Xanthomonas* species to induce hypertrophy of plant mesophyll cells (7), LntA, secreted by *Listeria monocytogenes* that stimulates a type III interferon response by inhibiting the heterochromatin-associated factor BAHD1, and Ank proteins secreted by *Orientia tsutsugamushi*, which downregulate transcription of host genes involved with histone modification and chromatin organization (8).

*Mycobacterium tuberculosis* (Mtb), the causative agent of tuberculosis (TB), is one of the leading causes of mortality worldwide. When Mtb is internalized by innate immune cells like macrophages, it establishes a replicative niche in phagosomes by manipulating a variety of defense pathways (lysosomal fusion, cell death, autophagy) (9–11). While traditionally thought of as a vacuolar pathogen, we now appreciate that cytosolic access, which results from ESX-1-dependent destabilization of the phagosome, is a major determinant of Mtb virulence (12, 13).

We also know, through mass spectrometry studies, that Mtb secretes upwards of 100 proteins into culture supernatants (14). Following phagosome destabilization, these proteins are predicted to access the macrophage cytosol and organelles. Supporting this prediction are protein-protein interaction studies that identified host factors that physically interact with Mtb secreted proteins and linked these interactions to specific Mtb infection outcomes (15–17).

Notable among these was a global IP-MS study from Penn et al, that reported high-confidence interactions between Mtb secreted proteins and nuclear host proteins involved in pre-mRNA splicing, mRNA export, and histone remodeling (15), suggesting that Mtb encodes and secretes proteins that can act as nucleomodulins. Supporting this idea, the Mtb secreted protein PtpA was recently shown to bind DNA, alter transcription, and promote immune suppression in a nuclear-import dependent fashion (18). Mtb also encodes several proteins capable of modifying host histones: Rv1988, which can methylate H3K42 (19), and Rv2067c, which can trimethylate H3K79 (20). Expression of each of these Mtb proteins has been linked to changes in host gene expression that support Mtb virulence.

Motivated by these foundational studies, we set out to catalog putative Mtb nucleomodulins and investigate their role in balancing the Mtb-macrophage host pathogen interface. Here, we report that Rv1075c, a previously described GDSL-like esterase (21), robustly accumulates in macrophage nuclei where it promotes expression of *Ifnb1* and select pro-inflammatory genes.

We also discovered a role for Rv1075c in modulating macrophage lipid metabolism, through altering the expression of lipid-associated genes, promoting lipid droplet formation, and helping sustain foamy macrophage populations in the lungs of infected mice. Our data suggest that Rv1075c achieves this regulation by interacting with and disrupting the SET1 histone methyltransferase complex. Together, these data describe a “moonlighting” role for this Mtb esterase in manipulating macrophage gene expression to promote bacterial persistence.

## RESULTS

### *M. tuberculosis* protein Rv1075c accumulates in macrophage nuclei

To identify Mtb nucleomodulins, we utilized a program called cNLS mapper that predicts importin α-dependent nuclear localization signals (NLS) (22, 23). According to the cNLS mapper system, a score of 8 or higher indicates exclusive localization to the nucleus, 6-8 indicates partial localization to the nucleus, 3-5 indicates localization to the cytoplasm and the nucleus, and a score of less than 3 indicates exclusive cytoplasmic localization. Because we were most interested in proteins with potential to be secreted from Mtb into host cells, we input a list of proteins previously identified in Mtb culture supernatants (14). Overall, 32/105 Mtb proteins analyzed received an NLS score >3.0, suggesting some likelihood of nuclear localization (**Fig. S1A**). The top 10 NLS scoring proteins are shown in **Fig 1A**. Of these, Rv1075c (NLS = 4.0) was notable because it was the only candidate with a predicted classical monopartite motif K(K/R)X(K/R), more specifically a class 1 classical nuclear localization signal KRIRL at amino acids 108-111.

**Figure 1:**
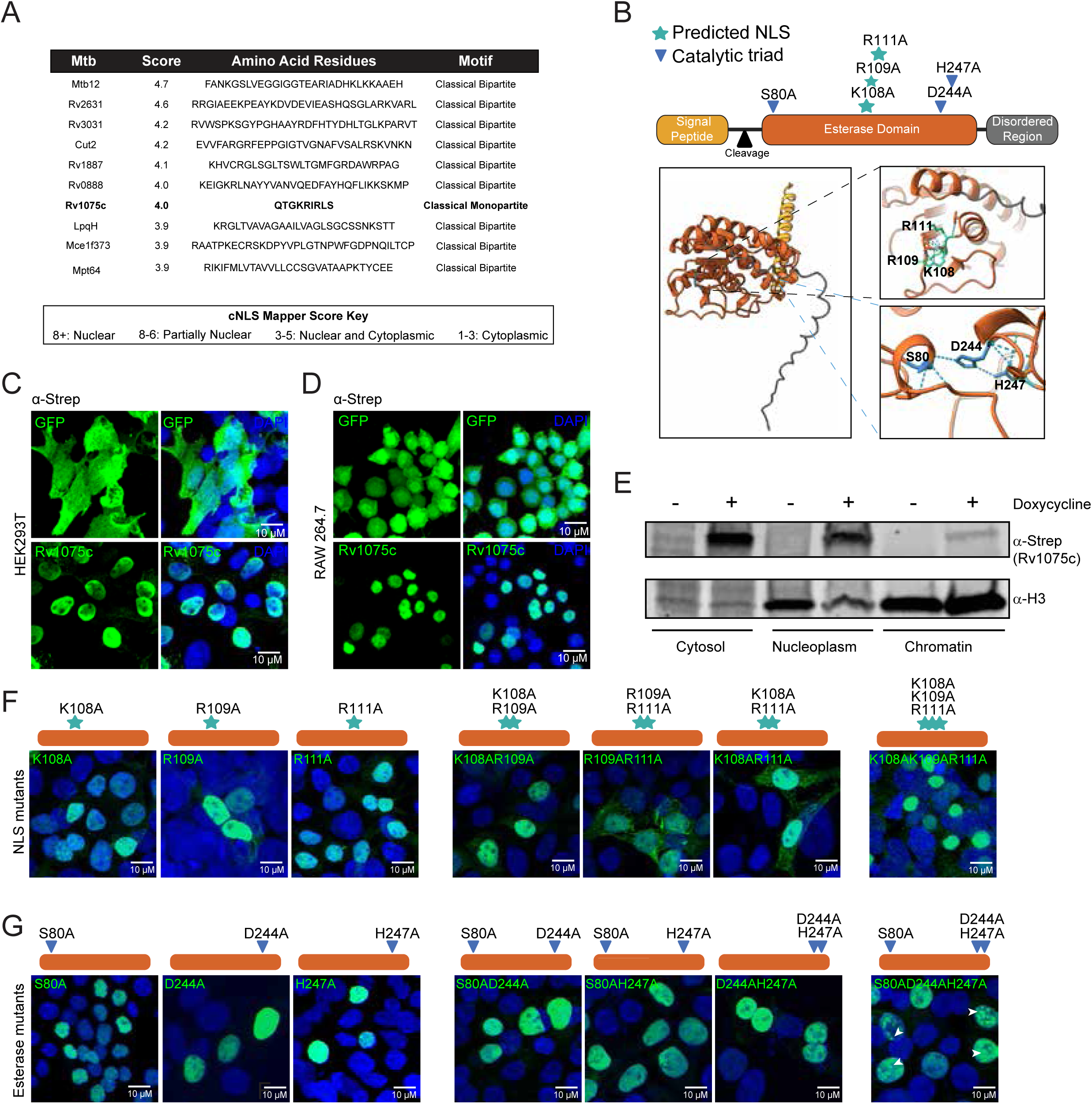
*M. tuberculosis* protein Rv1075c accumulates in macrophage nuclei. A. Table of putative nucleomodulins from Mtb with NLS score assigned by cNLS mapper B. Domain schematic and alpha fold prediction of Rv1075c. The cleaved signal peptide is labeled as yellow, the esterase domain is labeled as orange, and the disordered region is labeled as gray. The putative NLS amino acids are labeled as teal and denoted with a star. The catalytic triad amino acids are labeled as blue and denoted with a triangle. C. Immunofluorescence microscopy of HEK293T cells ectopically expressing C-2xSTREP GFP or C-2xSTREP Rv1075c. D. Immunofluorescence microscopy of doxycycline-inducible RAW 264.7 macrophages expressing C-2xSTREP GFP or C-2xSTREP Rv1075c. E. Immunoblot of doxycycline-inducible RAW 264.7 macrophages expressing C-2xSTREP Rv1075c biochemically fractionated into cytoplasm, nucleoplasm, and chromatin. F. Immunofluorescence microscopy of HEK293T cells expressing point mutations of the Rv1075c catalytic triad amino acids S^80A^, D^244A^, H^247A^. G. Immunofluorescence microscopy of HEK293T cells expressing point mutations of the Rv1075c putative NLS amino acids K^108A^, R^109A^, R^111A^.

Rv1075c is a conserved mycobacterial protein (**Fig, S1B**) reported to have esterase activity controlled by a catalytic triad of S80-D244-H247 (21). It is predicted to be translocated in a folded state out of Mtb by the twin arginine secretion pathway (24) (**Fig. 1B**). The predicted NLS KRIRL is conserved between *M. tuberculosis H37Rv, M. tuberculosis MTBC0,* and *M. bovis AF2122-97*, however *M. smegmatis* contains a glutamine instead of a lysine at position 108 and *M. marinum* has a valine in place of an isoleucine at position 110. Rv1075c was previously suggested to be required for Mtb survival in macrophages and can act as an esterase, with preference for short-chain fatty acids, particularly acetate (21). To determine the subcellular localization of Rv1075c in macrophages, we ectopically expressed a C-terminal 2xSTREP tagged version in HEK293T cells (**Fig. 1C**) and a tetracycline-inducible C-terminal 2xSTREP tagged version in RAW 264.7 macrophages (**Fig. 1D**). Strong nuclear localization was confirmed in both cell types by immunofluorescence microscopy (**Fig. 1C-D**). Chromatin association of 2xSTREP-Rv1075c was confirmed by immunoblot following biochemical fractionation of macrophages into cytosol, nucleoplasm, and chromatin-associated fractions (**Fig. 1E**).

To determine the role of Rv1075c’s putative NLS in the protein’s nuclear enrichment, we used site directed mutagenesis to sequentially create single, double, and triple mutants of the KRIRL^108-111^ sequence and transfected these constructs into HEK293T cells. We found that the single (K^108A^, R^109A^, R^111A^), double (K^108A^R^109A^, R^109A^R^111A^, K^108A^R^111A^), and triple mutants (K^108A^K^109A^R^111A^) of Rv1075c maintained strong nuclear localization, although two of the double mutants showed some cytosolic enrichment (R^109A^R^111A^, K^108A^R^111A^) (**Fig. 1F**). We then asked if the protein’s esterase activity, dictated by the S^80^-D^244^-H^247^ catalytic triad influenced the protein’s subcellular localization. When expressed in HEK293T cells, the esterase mutants accumulated in nuclei similarly to the wild-type control (**Fig. 1G**). Curiously, the triple mutant construct uniquely accumulated in puncta suggestive of a nuclear compartment (white arrows). Together, these data demonstrate that Mtb Rv1075c accumulates in mammalian cell nuclei independently of its putative NLS or its esterase enzymatic activity.

### *ΔRv1075c*-infected macrophages fail to fully induce *Ifnb1* and inflammatory genes

Because Rv1075c accumulates in macrophage nuclei, we posited that plays a role in manipulating host gene expression. To gain insight into how Rv1075c influences Mtb infection outcomes, we generated a ΔRv1075c Mtb strain using the oligonucleotide-mediated recombineering followed by Bxb1 integrase targeting (ORBIT) system (25). We confirmed generation of the KO strain by designing flanking PCR primers to amplify either a 1.2 kb band (WT Rv1075c) or a 3.2 kb band (KO Rv1075c). We also confirmed proper orientation of the cassette using primers to amplify the OriE and HygR regions within the payload plasmid (**Fig. S2A-B**). Compared to the parental pKM444 recombineering plasmid-containing ORBIT strain (parent), ΔRv1075c Mtb did not exhibit any defects replicating in culture (7H9 media supplemented with OADC) (**Fig. S2C**) nor did it display altered replication in bone marrow derived murine macrophages (BMDMs) (**Fig. 2A**). Additionally, ΔRv1075c infection (MOI=5) elicited the same cell death kinetics over a 16h time course as the parental control strain, as measured by propidium iodide incorporation (**Fig. 2B**).

**Figure 2:**
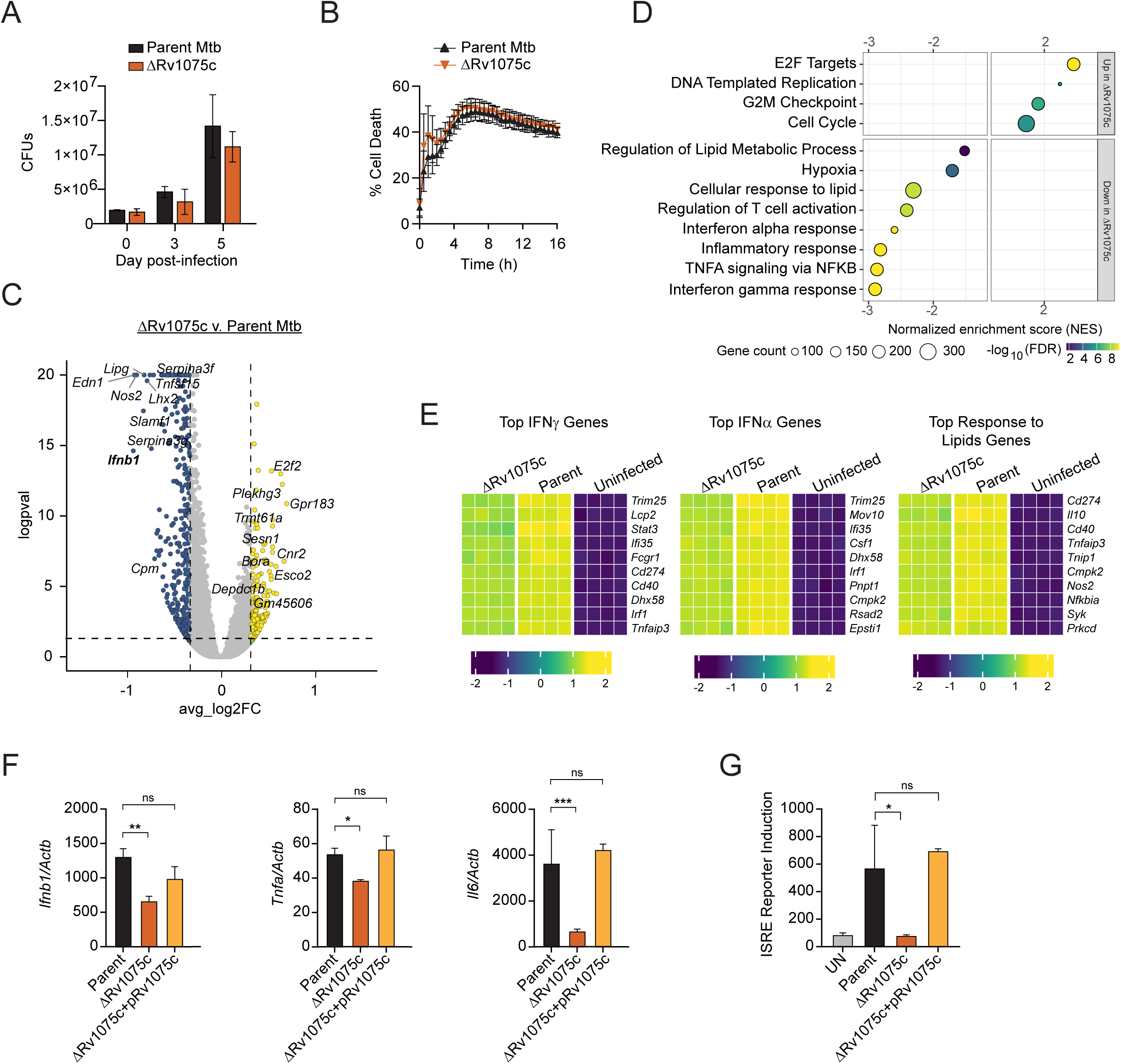
*ΔRv1075c*-infected macrophages fail to fully induce *Ifnb1* and inflammatory genes. A. Colony-forming units (CFU) quantification of Parent wild-type (Black) and ΔRv1075c (Orange) strains in BMDMs over five days (MOI 5). B. Propidium iodine incorporation over a 16h infection of BMDMs with Parent (Black) and ΔRv1075c (orange) Mtb strains (MOI 5). % = PI+ cells/total cells ∗ 100. C. Volcano plot of differentially expressed genes in ΔRv1075c versus Parent Mtb infected BMDMs 6 hours post infection. Upregulated genes (p<0.05; log2FC >0.5) in yellow and downregulated genes (p<0.05; log2FC >-0.5) in blue. D. Gene pathway enrichment analysis plot of up-and downregulated genes in ΔRv1075c versus Parent Mtb infected BMDMs for 6h. E. Heatmap of genes differentially expressed in ΔRv1075c vs Parent Mtb infected BMDMs in IFNγ, IFNα, or Response to Lipid categories. F. RT-qPCR of *Ifnb1*, *Tnfa*, and *Il6* in RAW 264.7 macrophages 6h post-infection with Parent, ΔRv1075c, or ΔRv1075c+pRv1075c Mtb relative to uninfected. One-way Anova. Error bars represent STDEV. **p<0.005. G. Secreted IFNβ in the supernatant of BMDMs (relative light units from ISRE reporter cells) infected with Parent, ΔRv1075c, or ΔRv1075c+pRv1075c Mtb 6 hours post infection. One-way Anova. Error bars represent STDEV. *p<0.05.

Having confirmed that infection with ΔRv1075c does not elicit significant differences in bacterial burden or cell death, we set out to understand how the absence of this nucleomodulin alters the Mtb-macrophage host pathogen interface. To this end, we infected BMDMs with parent or ΔRv1075c Mtb (MOI=5) for 6 hours and performed Illumina RNA-seq (PE 150, polyA+ selection). We found a total of 588 differentially expressed genes (DEGs) in BMDMs infected with ΔRv1075c vs. parent Mtb with 214 upregulated and 374 downregulated (**Fig. 2C**). Notably, the most downregulated gene was *Ifnb1*, which encodes IFN-β, a potent inducer of the type I IFN response in macrophages through autocrine and paracrine signaling manzanillo (13, 26).

Several downstream ISGs (*Rsad2*, *Il6*, *Slamf1, Ifi202b*) were also downregulated in the absence of Rv1075c (**Fig. 2C**). Pathway analysis highlighted downregulation of genes in the interferon alpha and gamma responses (the regulons of which are largely shared), as well as genes involved in canonical inflammation (*TNFA signaling via NFKB*) (**Fig. 2D-E, S2D**). We also noted downregulation of genes involved in lipid metabolism (**Fig. 2D-E, S2D**), an increasingly appreciated aspect of the macrophage response to Mtb (27–30). Upregulation of genes in the E2F transcription factor family and additional cell cycle-associated genes was also seen in ΔRv1075c Mtb-infected BMDMs (**Fig. S2E-F**). To confirm these gene expression changes, and more definitively ascribe them to loss of Rv1075c, we performed RT-qPCR of RAW 264.7 macrophages infected with our parent ORBIT strain, ΔRv1075c, or a complemented strain, which we generated by cloning Rv1075c in pMV306 a site-specific integrating expression plasmid containing the mycobacterium optimized promoter (MOPS) (**Fig. S2G-H**). Consistent with our RNA-seq, we found dampened expression of *Ifnb1*, *Tnfa*, and *Il6*, that was rescued by our complementation plasmid (**Fig. 2F**). Importantly, downregulation of *Ifnb1* at the transcript level was borne out at the level of protein secretion, as measured by ISRE reporter cells and supernatants of parent, ΔRv1075c, and ΔRv1075c + pRv1075c (**Fig. 2G**).

If Rv1075c directly mediates these gene expression changes, we hypothesized that its overexpression would elicit hyperinduction of the same genes. To test this, we employed a doxycycline-inducible Rv1075c RAW 264.7 cell line (**Fig. 1D** and **3A**). Because we did not detect any differences in gene expression in resting RAW 264.7 cells +/-Rv1075c (**Fig. 3B**), we supposed that macrophage activation, via infection or pattern recognition receptor engagement, was needed to turn on transcription of these genes. Cytosolic sensing of dsDNA through cGAS/STING plays a key role in activating *Ifnb1* and the type I IFN response during Mtb infection (12, 31, 32). Therefore, we transfected Rv1075c-expressing RAW 264.7 cells, alongside tetracycline-inducible GFP-expressing controls, with ISD and measured expression of Rv1075c-sensitive genes by RT-qPCR. We measured dramatic hyperinduction of *Ifnb1*, alongside several ISGs, at 6h post-ISD transfection (following 16h of doxycycline induction) (**Fig. 3C-D**). We observed a similar hyperinduction of *Ifnb1* and other inflammatory genes when Rv1075c-expressing cells were infected with Mtb (**Fig. 3E**). To investigate the contribution of Rv1075c’s enzymatic activity to this phenotype, we induced expression of GFP, wild-type Rv1075c, and an esterase mutant (S^80A-^D^244A-^H^247A^ = SDH) in tetracycline-inducible RAW 264.7 cells, transfected them with ISD as above, and measured expression of Rv1075c-sensitive genes by RT-qPCR. Consistent with a role for Rv1075c’s SDH domain in its nucleomodulin activity, the SDH mutant failed to hyperinduce *Ifnb1*, *Il6*, or *Rsad2*, resulting in expression levels similar to those of the GFP control (Fig. 2K). (**Fig 2K**). Together, these data demonstrate that Rv1075c is necessary and sufficient for maximal induction of *Ifnb1* and a subset of ISGs/inflammatory genes during Mtb infection and begin to implicate its esterase domain in its gene regulatory activity.

**Figure 3:**
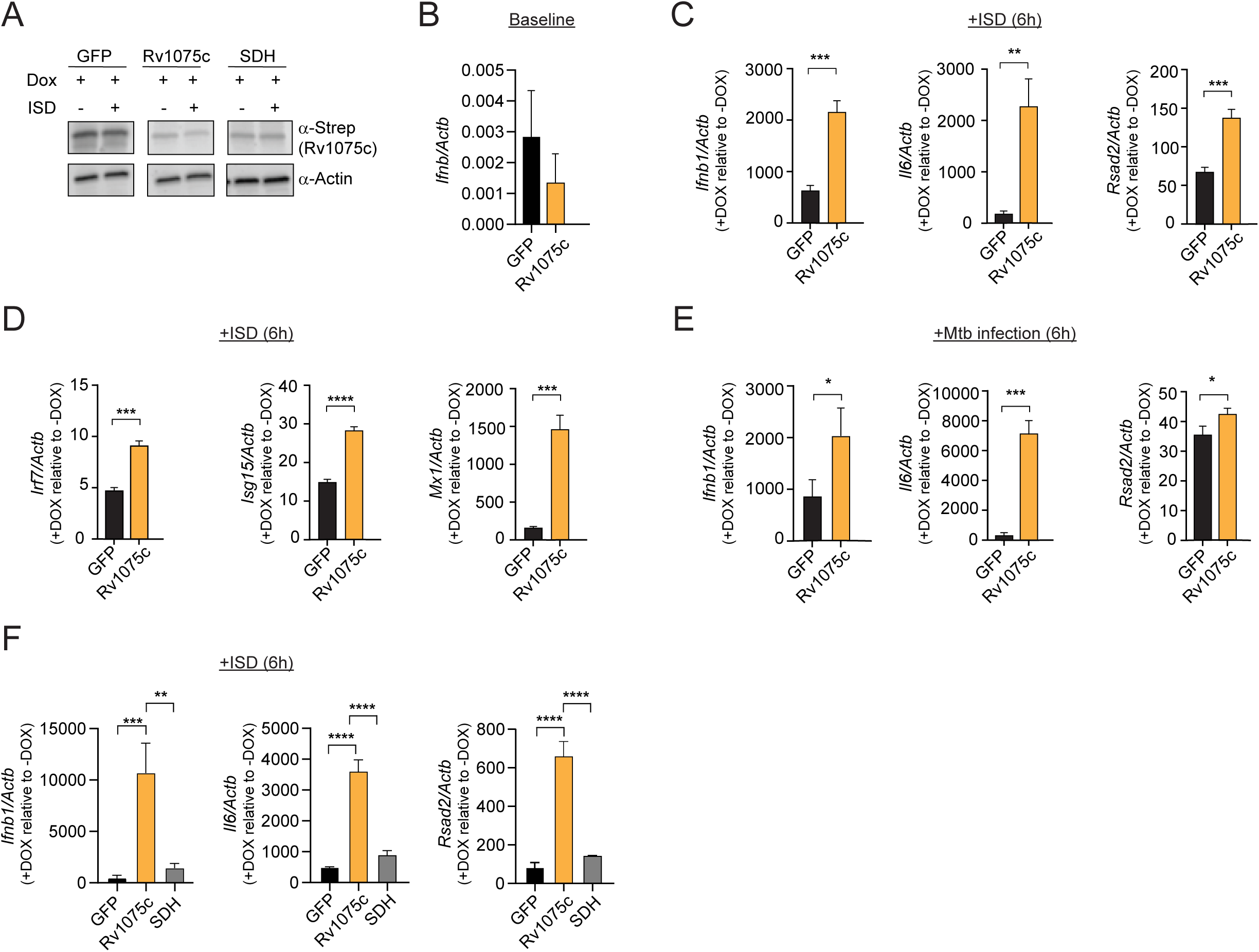
Rv1075c overexpression promotes hyperinduction of *Ifnb1* and inflammatory genes in activated macrophages. A. Immunoblot of C-2xStrep-Rv1075c overexpression in the absence and presence of doxycycline (anti-Strep). B. RT-qPCR of transcript *Ifnb1* in doxycycline-inducible RAW 264.7 cells expressing either GFP or Rv1075c for 16h. Student T test used. ***p<0.0005, ****p<0.0001. C. RT-qPCR of *Ifnb1*, *Il6*, and *Rsad2* in Rv1075c (orange) or GFP (black)-overexpressing 6h post-ISD transfection, relative to doxyclinetrated. Student T-test. Error bars represent STDEV. ***p<0.0005. D. As in C but for *Irf7*, *Rsad2*, *Isg15*, and *Mx1*. E. As in C but 6h post-Mtb (wild-type Erdman) infection (MOI=5). Student T-test. Error bars represent STDEV. *p<0.05, ***p<0.0005. F. RT-qPCR of *Ifnb1*, *Il6*, and *Rsad2* in Rv1075c (orange), S80AD244AH247A (gray), or GFP (black)-overexpressing 6h post-ISD transfection, relative to doxycline. One way Anova. Error bars represent STDEV. ** p<0.005, **** p<0.0001.

### Rv1075c manipulates host lipid metabolism in macrophages

In addition to defects inducing *Ifnb1* and inflammatory genes, we also noticed dampened expression of lipid metabolism genes in ΔRv1075c-infected macrophages (**Fig. 2D-E**).

Regulation of lipid metabolism is tightly linked to macrophage control of Mtb and lipid-laden macrophages are known to support Mtb survival and replication (29, 30). To further explore a potential link between Rv1075c and lipid gene expression, we infected RAW 264.7 macrophages with parent, ΔRv1075c, or ΔRv1075c + pRv1075c strains (MOI=5) and measured expression of several lipid genes at 6h post-infection. Consistent with our RNA-seq, ΔRv1075c-infected macrophages had lower expression of *Hilpda*, a positive regulator of lipid droplet formation and *Sphk1*, an enzyme that converts sphingosine into sphingosine-1-phosphate (**Fig. 4A**). Rv1075c overexpression, on the other hand, promoted modest upregulation of *Hilpda* and hyperinduction of the lipase *Lipg* in Mtb-infected RAW 264.7 cells (**Fig. 4B**), supporting a role for Rv1075c in modulating the expression of select lipid genes.

**Figure 4.**
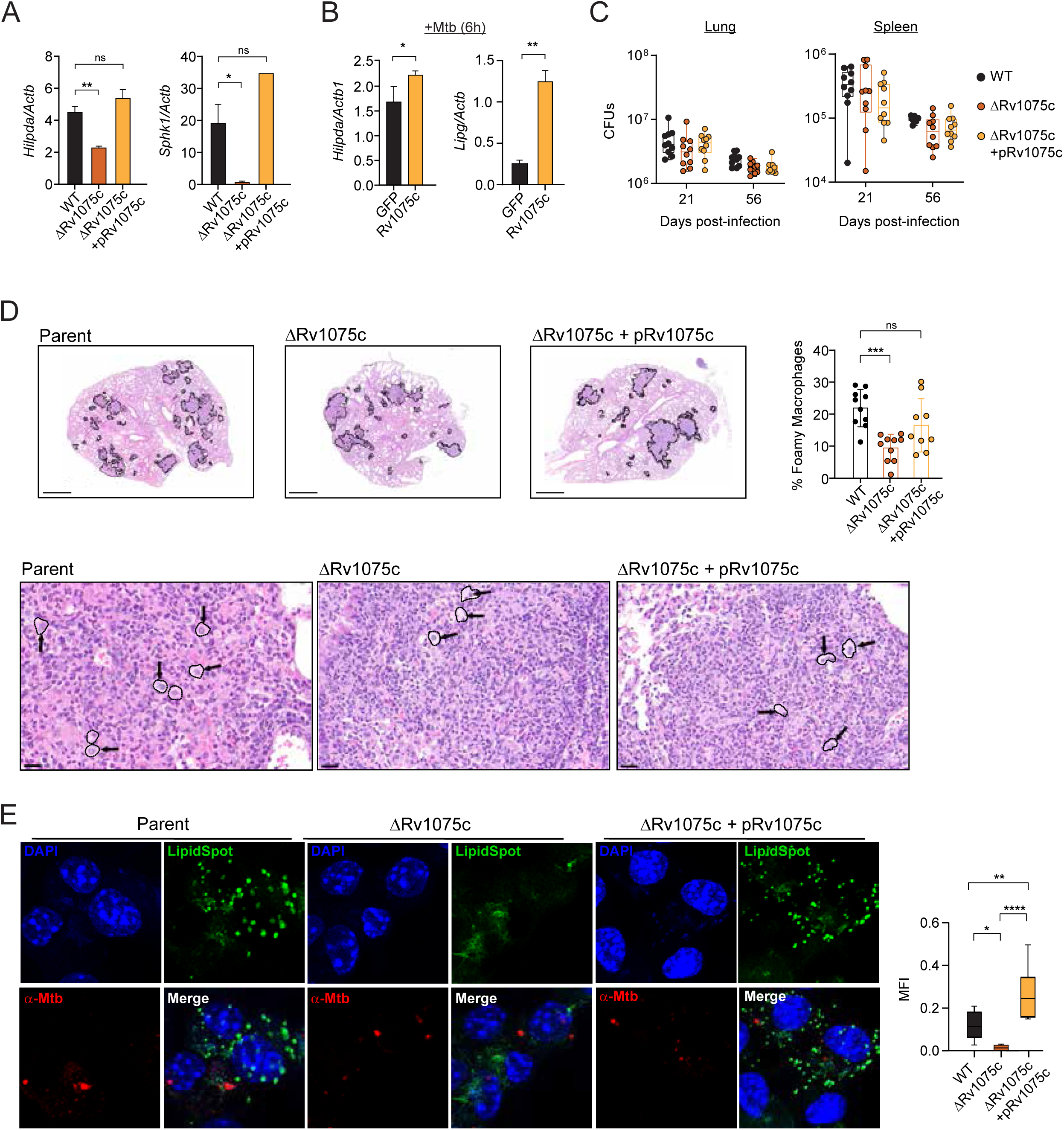
Rv1075c manipulates host lipid metabolism in macrophages. A. RT-qPCR of *Hilpda* and *Sphk1* in RAW 264.7 macrophages 6h post-infection with Parent (black), ΔRv1075c (dark orange), or ΔRv1075c+pRv1075c (light orange) Mtb (MOI=5). One-way Anova. Error bars represent STDEV. *p<0.05, **p<0.005. B. As in A, but for *Hilpda* and *LipG* in doxycycline-inducible RAW 264.7 macrophages expressing Rv1075c or GFP 6h post-Mtb infection (wildtype Erdman, MOI=5). One way Anova. Error bars represent STDEV. *p<0.05, **p<0.005. C. Colony-forming units (CFUs) recovered from the lung and spleen of mice infected with Parent (black), ΔRv1075c (dark orange), or ΔRv1075c+pRv1075c (light orange) at Day 21 and 56 post-infection. D. Hematoxylin and eosin (H&E) stain of regions of inflammation in the lungs of mice infected with Parent, ÄRv1075c, or ÄRv1075c+pRv1075c Mtb at Day 21. Within these regions, foamy macrophages were scored by a veterinary pathologist and are denoted by black arrows and circles. One-way Anova. Error bars represent STDEV. ***p<0.0005. E. LipidSpot (green), nuclei (blue, DAPI), and Mtb (red, anti-Mtb) IF microscopy in Parent, ΔRv1075c, and ΔRv1075c+pRv1075c infected BMDMs. Quantification of LipidSpot MFI in infected BMDMs. One-Way ANOVA. Error bars represent STDEV. *p< 0.05.

Having implicated Rv1075c in the macrophage transcriptional response to Mtb (**Fig. 2-3**), we sought to determine a role for this protein during *in vivo* infection. To this end we performed a low dose aerosol infection (∼100 CFUs) of wild-type C57BL6 mice with parent, ΔRv1075c, or ΔRv1075c + pRv1075c strains. At day 21 and 56 post-infection, we measured no significant differences in CFUs recovered from the lung or spleen between the three genotypes, although ΔRv1075c CFUs trended downward (**Fig. 4C**). Although several gross measures of lung pathology, including inflammation and the percentage of grids containing neutrophils, were similar among mice infected with the three strains (**Fig. S3A**), we observed significantly fewer foamy macrophages in the lungs of mice infected with ΔRv1075c than in those infected with either the parent or complemented strains (**Fig. 4D**). Motivated to further investigate Rv1075c’s role in manipulating macrophage lipid metabolism, we infected BMDMs with parent, ΔRv1075c, or ΔRv1075c + pRv1075c strains and visualized lipid droplets (green, LipidSpot) and Mtb (red, anti-Mtb) via immunofluorescence microscopy. Consistent with failure to elicit foamy macrophages in the lungs of infected mice, ΔRv1075c simulated significantly fewer lipid droplets during BMDM infection (**Fig. 4E**). Collectively, these findings support a role for Rv1075c in reprogramming macrophage lipid metabolism and promoting the formation of lipid-rich cellular niches during Mtb infection.

### Rv1075c interacts with the host SET1 methyltransferase complex

To gain insight into the molecular mechanisms underlying Rv1075c’s ability to manipulate inflammatory and lipid gene expression in macrophages, we next sought to identify Rv1075c-interacting proteins. Briefly, we ectopically expressed 2xStrep-Rv1075c or 2xStrep-GFP in HEK293T cells and performed immunoprecipitation from nuclear fractions followed by mass spectrometry. We identified hundreds of proteins that were enriched in our Rv1075c pulldowns compared to GFP (**Table S1**). At the top of this list were many nuclear proteins and several mitochondrial proteins. Because we do not see ectopically-expressed Rv1075c localize to mitochondria via immunofluorescence microscopy, these mitochondrial interactors likely result from mitochondrial membrane contaminants in our nuclear-enriched pellets. Similarly, because Rv1075c appears to be excluded from the nucleolus in our IF images (**Fig. S4A**), we chose not to pursue interactions between Rv1075c and several abundant nucleolar proteins. Instead, we focused on attention on interactions between Rv1075c and components of the SET1/COMPASS histone methyltransferase complex: RBBP5, WDR5, and ASH2L (**Fig. 5A**). SET1 is a highly conserved, multi-subunit enzyme complex that catalyzes the methylation of histone H3 at lysine 4 (H3K4). SET1-directed deposition of H3K4me3 has been implicated in cell cycle regulation and DNA repair (33–35), but its role in controlling expression of innate immune genes is less well-understood. To validate our mass-spectrometry data, we immunopurified Rv1075c from tetracycline-inducible RAW 264.7 cells, alongside a GFP control, and confirmed Rv1075c-specific pulldown of ASH2L, RBBP5, and WDR5 using antibodies directed against the endogenous proteins (**Fig. 5B**). We were also able to pulldown another member of the complex, DPY30, despite not picking it up via mass-spectrometry (**Fig. 5B**). Immunofluorescence microscopy of each of the SET1 complex proteins confirmed that like Rv1075c, they are predominantly nuclear and exhibit a diffuse staining pattern (**Fig. 5C**). The interaction between SET1 proteins and Rv1075c appeared to be independent of the protein’s esterase activity, as the D244A-H247A and S80A-D244A-H247A mutants were still able to immunoprecipitate ASH2L and RBBP5. If anything, the interaction between the Rv1075c esterase mutants and SET1 components was stronger than that observed with wild-type Rv1075c, suggesting that the esterase domain may play a role in dissociating Rv1075c from the SET1 complex. (**Fig. S4B**).

**Figure 5.**
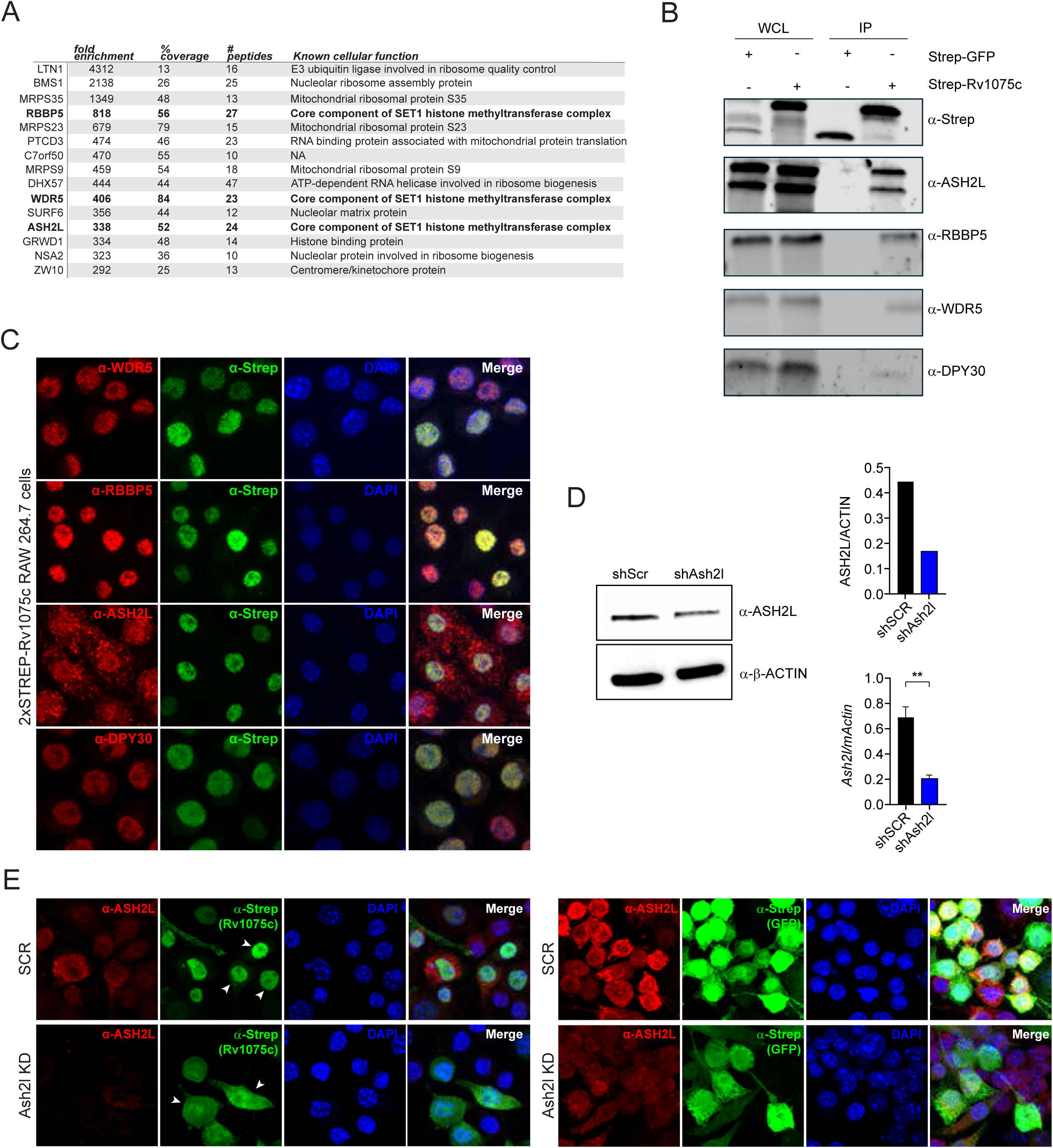
Rv1075c interacts with the host SET1 methyltransferase complex. A. Results of IP-Mass spectrometry of C-2xSTREP-Rv1075c and C-2xSTREP-GFP expressed in HEK293T cells. Interacting proteins ranked by fold enrichment (Rv1075c over GFP) B. Immunoblot of Strep-IP of Rv1075c, probed for endogenous Ash2l (anti-Ash2l), Wdr5 (anti-Wdr5), Rbbp5 (anti-Rbbp5), and Dpy30 (anti-Dpy30) from HEK 293T cells. C. IF microscopy of doxycycline-inducible RAW 264.7 macrophages expressing Rv1075c (green, anti-Strep) and Wdr5, Rbbp5, Ash2l, or Dpy30 (red, anti-SET1 protein). D. Protein and transcript expression of Ash2L (anti-Ash2l) in shScr and shAsh2l RAW 264.7 macrophages. Student T test. Error bars represent STDEV. **p<0.005 E. IF microscopy of doxycycline-inducible RAW 264.7 macrophages expressing C-2xSTREP-Rv1075c or C-2xSTREP-GFP (green, anti-Strep) and transduced with shRNA construct targeting Ash2l, alongside a SCR control (red, anti-Ash2l). Arrows denote Rv1075c localization.

Remembering our earlier failure to disrupt Rv1075c nuclear accumulation via mutation of its putative NLS (**Fig. 1F**), we hypothesized that Rv1075c gets to the macrophage nucleus through its interaction with SET1 components (i.e. a piggyback model of nuclear translocation). To test this, we introduced an shRNA knockdown construct directed against *Ash2l* via lentiviral transduction of RAW 264.7 cell lines expressing 2xSTREP-Rv1075c or 2xSTREP-GFP and verified knockdown of ASH2L at the protein and RNA level (**Fig. 5D**). Via immunofluorescence microscopy, we observed striking redistribution of Rv1075c from nuclei to the cytosol in macrophages lacking ASH2L (**Fig. 5E**), suggesting Rv1075c piggybacks on SET1 proteins (likely newly translated ones) to translocate into the macrophage nucleus.

### SET1 complex knockdown hyperinduces inflammatory and lipid gene expression in activated macrophages

Because our data suggest that Rv1075c interacts with SET1 components to modulate macrophage gene expression during Mtb infection, we hypothesized that loss of SET1 would phenocopy Rv1075c expression. To test this, we took our *Ash2l* KD cell lines, alongside *Rbbp5* KD cell lines (**Fig. S5A**), stimulated them with ISD or infected them with Mtb, and measured expression of previously identified Rv1075c-sensitive genes. Consistent with SET1 controlling macrophage gene expression during Mtb infection, we found that loss of *Ash2l* and *Rbbp5* resulted in hyperinduction of inflammatory (**Fig. 6A-B**) and lipid metabolism (**Fig. 6C-D**) genes induced as part of the response to ISD. In response to Mtb infection, SET1 KDs similarly hyperinduced inflammatory genes (**Fig. 6E-F**) but interestingly, showed the opposite phenotype for lipid metabolism genes (i.e. SET1 KDs had lower expression of *Hilpda* and no difference in *Spnk1* expression compared to SCR controls (**Fig. S5B-C**)), suggesting a more complex regulatory network for these genes in the context of infection. Collectively, these data are consistent with a model whereby SET1 deposition of H3K4me3 dampens induction of inflammatory genes during macrophage activation, and Rv1075c expression disrupts SET1 activity. Consistent with this, pre-treatment of BMDMs with OICR 9429, a steric inhibitor of the SET1 complex (36), also led to hyperinduction of *Ifnb1*, *Rsad2*, and *Il6* (**Fig. 6G**), phenocopying expression of Rv1075c.

**Figure 6.**
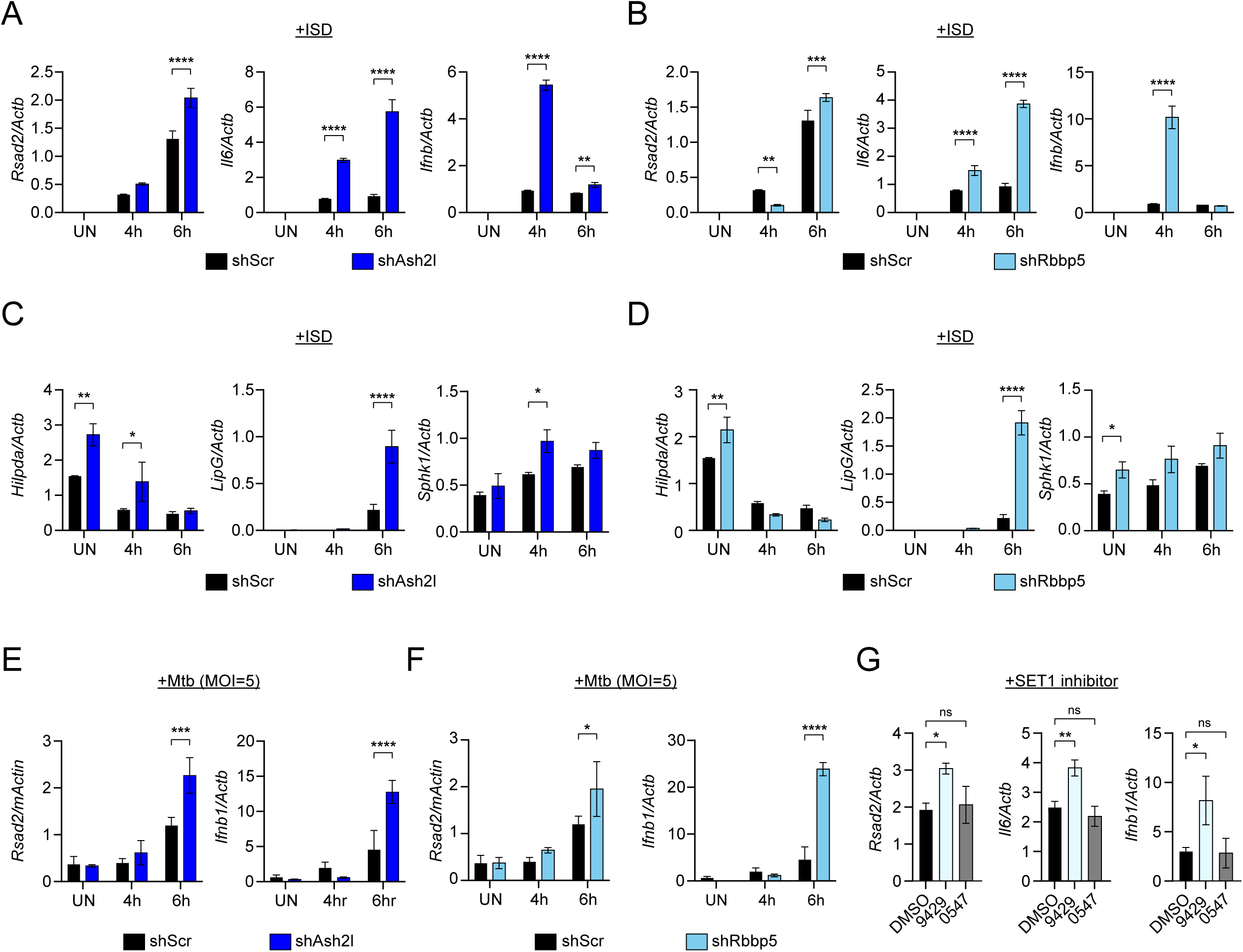
Loss of SET1 complex members elicits hyperinflammation in activated macrophages. A. RT-qPCR of *Rsad2*, *Il6,* and *Ifnb1* in shAsh2l (dark blue) and shScr (black) RAW 264.7 macrophages transfected with ISD for 4 and 6 hours. Two-way Anova. Error bars represent STDEV. **p<.005, ****p <0.0001. B. As in A, but for shRbbp5 (light blue) and shScr (black) RAW 264.7 macrophages. C. As in A, but for *Hilpda* and *LipG* in shAsh2l (dark blue) and shScr (black) RAW 264.7 macrophages. D. As in A, but for *Hilpda* and *LipG* in shRbbp5 (light blue) and shScr (black) RAW 264.7 macrophages E. RT-qPCR of *Rsad2* and *Ifnb1* in shAsh2l (dark blue) and shScr (black) RAW 264.7 macrophages infected with Mtb (MOI=5) for 4 and 6 hours. Two-way Anova. Error bars represent STDEV. *p<0.05, ***p<0.0005. F. As in E, but for *Rsad2* and *Ifnb1* in shAsh2l (light blue) and shScr (black) RAW 264.7 macrophages G. RT-qPCR of *Rsad2*, *Il6,* and *Ifnb1* in BMDMs treated overnight with DMSO (black), the SET1 methyltransferase inhibitor OICR 9429 (10 uM, light gray), and the inhibitor control OICR 0547 (10 uM, dark gray) then infected with Mtb (wild-type Erdman, MOI=5) for 6 hours. One-way Anova. Error bars represent STDEV. *p<0.05, **p<0.005.

## DISCUSSION

The success of Mtb as a pathogen stems largely from its ability to establish an immune milieu that supports its persistence, replication, and spread. Here we demonstrate that Rv1075c, a previously described Mtb secreted protein with esterase activity, uses the host cell SET1 histone methyltransferase complex to translocate into macrophage nuclei and tune the expression of select inflammatory and lipid genes. Our findings suggest that Mtb uses Rv1075c create a pro-bacterial niche in macrophages by polarizing macrophages towards a type I IFN response and promoting foamy macrophage formation. While the precise molecular mechanisms underpinning these phenotypes remain unclear, our data suggest that these phenotypes are driven, at least in part, via inhibition of the SET1 complex (**Fig. 6**).

SET1 and its associated histone mark H3K4me3 are largely associated with active transcription, with highest deposition of H3K4me3 seen around transcription start sites (37, 38). Under this paradigm, it seems counterintuitive that loss of SET1, via knockdown of *Ash2l* or *Rbbp5*, results in hyperinduction of *Ifnb1*, *Il6*, and *Rsad2* following macrophage activation (**Fig. 6**). One possible explanation for this paradoxical result is that cells compensate for loss of SET1 by altering other chromatin remodeling complexes (upregulating activating marks or downregulating inhibitory marks) resulting in a genome that is hyperresponsive to inflammatory cues. Another possibility is that inhibiting SET1 allows for deposition of another, more potent activating mark at innate immune genes. Recent work in *Candida albicans* demonstrated that deletion of SET1 allows for acetylation of H3K4 residues formerly marked by methylation, and that this switch from H3K4me3 to H3K4Ac enables a transient transcriptional burst at inducible genes (39). It is possible that by inhibiting the SET1 complex, Rv1075c promotes increased deposition of H3K4Ac at select inflammatory genes to promote Mtb pathogenesis. To formally test this hypothesis, one would want to perform a genome-wide analysis of H3K4me3 and H3K4Ac in cells +/-Rv1075c. We actually attempted a version of this, performing CUT&RUN for H3K4me3 in ISD-transfected macrophages +/-doxycycline (to induce expression of Rv1075c), but the pervasiveness of the H3K4me3 mark and the broad nature of its peaks made peak calling difficult, leaving us hesitant to draw major conclusions about how Rv1075c expression broadly impacts H3K4me3 levels. An alternative approach like ATAC-seq may provide more insight into how Rv1075c alters chromatin organization to promote expression of select innate immune genes during Mtb infection.

Another key outstanding question pertains to the mechanism through which Rv1075c regulates lipid droplets and foamy macrophage formation (**Fig. 4D-E**). Consistent with Rv1075c acting via its control of SET1, we do measure dysregulation of lipid-associated transcripts in *Ash2l* and *Rbbp5* KD cell lines (**Fig. 6C-D**). One Rv1075-sensitive lipid gene encodes HILPDA (Hypoxia-Inducible Lipid Droplet-Associated Protein), a protein with a well-established role in promoting the accumulation of triglycerides and promoting lipid storage (40, 41). Another Rv1075c-sensitive gene encodes LIPG, which can help trigger the accumulation of intracellular lipid droplets in response to oxidative stress (42). Thus, the downregulation of *Hilpda* and *Lipg* expression seen in ΔRv1075c infected macrophages could contribute to the lipid droplet phenotype reported (**Fig. 4E**). It is also possible that Rv1075c uses its esterase activity to modulate fatty acid pools in infected cells. Acetate, a preferred substrate of Rv1075c (21), is converted into acetyl-coA, which serves as a building block for fatty acid synthesis and triacylglycerol storage in lipid droplets. Rv1075c could influence lipid droplet formation by controlling the balance of acetate/acetyl-coA in macrophages. Experiments exploring how esterase mutant Rv1075c constructs impact lipid droplet phenotypes and/or the generation of esterase mutant-expressing Mtb will help directly test these non-mutually exclusive models.

Earlier studies of Rv1075c found that loss of Rv1075c in Mtb resulted in a half log decrease in recoverable CFUs at 28 and 56 days post-infection of C3HeB/FeJ mice (21). We did not observe a comparable reduction in bacterial burden in our C57BL/6 mouse model (**Fig. 3C**). This discrepancy could stem from the Mtb strain background (our highly-virulent Erdman strain vs. the less virulent CDC1551-background transposon mutant used in Yang et al. (21)). These differences might also be attributable to the mouse strains. Despite being a common pre-clinical model to study Mtb pathogenesis, C57BL6 mice are quite resistant to Mtb, stemming in part from their failure to mount robust type I IFN responses. The C3HeB/FeJ mice infected by Yang et al. may more accurately reflect the role of Rv1075c in Mtb pathogenesis, such that mice are better able to restrict Mtb without the extra boost of type I IFN stimulated by Rv1075c.

While this manuscript was in preparation, Chen et al. reported that Rv1075c interacts with the SET1 histone methyltransferase complex to regulate macrophage gene expression (43).

Although our studies share several similarities, our conclusions differ in important ways. First, Chen et al. identify two NLS motifs as essential for nuclear localization, whereas our mutants showed minimal effects on localization, potentially reflecting differences in mutagenesis strategy (deletion vs. alanine substitution). Second, they conclude that Rv1075c suppresses inflammatory gene expression, whereas our data support a pro-inflammatory role. This might stem from a difference in infection models (avirulent *M. bovis* vs. virulent Mtb), the timepoints examined (6h in our studies vs. 24h in Chen et al.), or perhaps most likely, the fact that avirulent *M. bovis* does not strongly activate type I IFN responses in macrophages. Finally, they report a significant pathogenesis defect for ΔRv1075c strains *in vivo*, following intratracheal installation of 10^7^ BCG bacilli into C57BL/6 mice. We maintain that our low dose aerosol infection with virulent Mtb provides a better representation of how Rv1075c may function in *bona fide* tuberculosis disease, with the caveat that we may be missing type I IFN-relevant phenotypes in the C57BL/6 mouse. Despite these experimental discrepancies, these studies collectively highlight a previously unrecognized chromatin-mediated axis in the Mtb-macrophage host-pathogen interface that governs inflammation and lipid metabolism.

## ACKNOWLEDGMENTS

We thank the members of the Watson and Patrick labs for their critical review and feedback in the preparation of this manuscript. Imaging was done with the help of Malea Murphy at the Integrated Microscopy and Imaging Laboratory at the Texas A&M Naresh K. Vashisht College of Medicine and the Vanderbilt Cell Imaging Shared Resource (supported by NIH grants CA68485, DK20593, DK58404, DK59637 and EY08126). Mass spectrometry of Rv1075c interacting proteins was carried out by the University of Texas Southwestern Medical Center Proteomics Core. Library preparation and RNA-sequencing was conducted by the Genomic and RNA Profiling Core at Baylor College of Medicine. This work was supported in part by R21AI203299 to KLP, R01AI155621 to ROW and KLP, and R01AI179037 to ROW and KLP. Trainees were supported by F31AI176652 to AKC, F31AI176795 to CJM, F31CA298557 to MHS, F31AI197784 to KSA, 5T32GM137793 to JBH, and T32GM135748 to AKC.

## Materials and Methods

### Cell culture

RAW 264.7 macrophages, HEK293T, L929 ISRE, and Lenti X cells were cultured at 37°C with 5% CO2. Cell culture medium was comprised of High glucose, sodium pyruvate, Dulbecco’s modified Eagle medium (Thermo Fisher) with heat inactivated 10% FBS (Gibco), and 0.5% HEPES (Cytiva). Bone marrow derived macrophages (BMDMs) were differentiated from bone marrow cells isolated by washing mouse femurs with 10 mL DMEM 1 mM sodium pyruvate.

Cells were centrifuged for 5 min at 400 rcf and resuspended in BMDM media (DMEM, 20% FBS (Gibco), 1 mM sodium pyruvate (Lonza), and 10% MCSF conditioned media. Bone marrow cells were counted and plated at 5×106 cells per 15 cm non-tissue culture treated dishes in 30 mL complete BMDM media. Cells were fed with an additional 15 mL of BMDM media on day 3.

Cells were harvested on day 7 with 1x PBS EDTA (Lonza). Lenti-X cells were used to produce lentiviral particles. Where necessary, RAW 264.7 cells were selected with and maintained in 5 μg/ml puromycin (InvivoGen), 5 μg/ml blasticidin (InvivoGen), and/or 100µg/mL hygromycin (Gibco, 10687010). For infections, antibiotics were omitted from culture media. Tagged expression constructs were made by first cloning cDNAs from RAW 264.7 RNA into pENTR1a entry vectors modified to contain in-frame epitope and constructs were verified utilizing Plasmidsarus. Constructs were then Gateway cloned with LR Clonase (Invitrogen) into pLenti destination vectors (Addgene, plasmid 19067). Expression of tagged proteins was confirmed by transfecting HEK293Ts with 1 μg of pDEST and harvesting cell lysates after 1 to 2 days of expression. Proteins were separated by SDS-PAGE and visualized by Western blotting with primary antibodies for STREP (genscript A01732-100and or endogenous antibodies (αRbbp5 cell signaling 13171s, αAsh2l - Abcam ab176334). To make RAW 264.7 stable expression cells lines, Lenti-X 293T cells were co-transfected with pLenti plasmids and the packaging plasmids psPAX2 and pMD2G/VSV-G (Addgene, plasmids 12259 and 12260) to produce lentiviral particles. Macrophage cell lines were transduced with lentivirus for two consecutive days plus according to the manufacturer’s instructions using a ratio of 3 μL Lipofectamine 2000 per 1 μg plasmid DNA (Invitrogen) and selected with antibiotic 48 hours post transfection. Expression of tagged proteins was confirmed by Western blotting with antibodies against corresponding epitope tags.

### Tetracycline inducible cell line generation

For generation of tetracycline inducible RAW 264.7 Macrophages, pLenti CMV rtTA3 Blast (Addgene w756-1) stably expressing clonal RAW 264.7 Macrophages were transduced with pLenti CMV Puro DEST (Addgene w118-1) constructs containing 2xStrep-GFP or 2xStrep-Rv1075c. After 48h, cells were selected through addition of puromycin (Invivogen). 1 mg/mL doxycycline (Sigma-Aldrich) treatment was used to activate construct expression.

### Stable shRNA Knockdown generation

For RAW 264.7 Macrophages stably expressing scramble knockdown and Srsf6 knockdown, lentivirus was made by transfection with a pSICO scramble non-targeting shRNA construct and pSICO *Ash2l (*targeting exon 3) *and Rbbp5* (targeting 3’ UTR) shRNA constructs using polyjet (SignaGen Laboratories, SL100688). Virus was collected 24 and 48 hours post transfection.

RAW 264.7 Macrophages were transduced using lipofectamine 2000 (Thermo Fischer, 52887). After 48 hours, media was supplemented with hygromycin (Invitrogen, 10687010) to select for cells containing the shRNA plasmid.

### Mycobacterium tuberculosis strains

The Erdman strain (Erdman WT, Erdman mCherry) was used for *M. tuberculosis* infections. Low passage lab stocks were thawed for each experiment to ensure virulence was preserved. M. tuberculosis was cultured in roller bottles at 37°C in Middlebrook 7H9 broth (BD Biosciences) supplemented with 10% OADC (BD Biosciences), 0.5% glycerol (Fisher), and 0.1% Tween-80 (Sigma) or on 7H10 plates. All work with *M. tuberculosis* was performed under Biosafety level 3 containment using procedures approved by the Texas A&M University and Vanderbilt University Medical Center Institutional Biosafety Committees.

### ORBIT Mtb Cloning

The ΔRv1075c mutant was made following the protocol of Muphy et al. Briefly, Mtb strains were electroporated with pKM444 and grown inn 30 ml 7H9 media containing OADC, 0.2% glycerol, 0.05% tween 80, and 20ug/ml kanamycin. At an OD of 0.8, 500 ng/mL of ATc was added to the culture. After 8 hours of incubation at 37°C, 3 ml of 2 M glycine was added to the culture. The cells were shaken at 37°C overnight (16 h to 20 h in total following induction), collected by centrifugation, the supernatant was removed, and the pellet was gently resuspended in 2 ml of 10% cold glycerol and then brought up to 20 ml with 10% cold glycerol. The centrifugation and washing steps were repeated. After the second wash, the cells were collected by centrifugation and resuspended in 2 ml of 10% cold glycerol. Aliquots (280 μl) of electrocompetent cells were added to sterile Eppendorf tubes containing 1 μg of an *attP*-containing oligonucleotide and 200 ng of the *attB*-containing plasmid cells and DNA were mixed by pipetting and transferred to ice-cooled electroporation cuvettes (0.2 cm path length). The cells were shocked with an electroporator at settings of 2.5 kV, 1,000 Ω, and 25 μF. Following electroporation, the cells were resuspended in 2 ml 7H9 media and rolled at 37°C overnight. Recombinant candidate colonies were picked after 3 weeks into 5 ml 7H9-OADC-Tween containing 50 μg/ml hygromycin and grown with shaking for 4 to 5 days at 37°C. Positive clones were verified by PCR.

### RNA sequencing

RNA-seq and analysis RNA-seq analysis was performed on BMDMs infected with WT, ΔRv1075c, or ΔRv1075c + pRv1075c after 6 hours. Bulk RNA-seq was performed in biological triplicate by the Baylor College of Medicine Genomic and RNA Profiling core. Reads were aligned to the Mus musculus reference genome build GRCm39 and analyzed using ROSALIND software (San Diego, CA), which uses a hyperscale architecture developed by ROSALIND, Inc. Differentially expressed genes were identified using a p-value threshold of <0.05. Volcano plots were generated in RStudio using raw counts.

### Extracellular IFN-β Luciferase Assay

Macrophage-secreted type I IFN-β levels were determined using a L929 cells stably expressing a luciferase reporter gene under the regulation of type I IFN signaling pathway (L929 ISRE cells). 25×10^6^ L929 ISRE cells were seeded in a clear 12-well plate and incubated at 37 °C with 5% CO2 the previous day. Macrophage supernatant was collected 6 hours post infection, diluted 1:2 in complete media and transferred to the L929 ISRE cells and incubated for 6 hr.

Cells were washed with PBS, lysed in 30 μL cell culture lysis buffer, and transferred to a white 96-well flat-bottomed plate (Costar, 3693). 30 μL of Luciferase Assay System substrate solution (Promega, E1501) was added to the plate and luminescence read immediately using a Cytation5 plate reader (Biotek).

### Mouse strains

Mouse husbandry and strain C57BL/6J mutator mice were purchased from the Jackson laboratory. All mice used in experiments were compared to age-and sex-matched controls and fed 4% standard chow. Littermate controls were used in all experiments. For ex vivo BMDM experiments, male mice between 10 and 12 weeks of age were used. For in vivo Mtb infections, mice were infected at 12 weeks of age.

### *In vitro* Bacterial Infections

Multiplicity of infection (MOI) for each bacterial infection is listed by experiment in the figure legends. The Erdman strain was used for all M. tuberculosis infections. Low-passage lab stocks were thawed for each experiment to ensure that virulence was preserved. M. tuberculosis was cultured in roller bottles at 37°C in Middlebrook 7H9 broth (BD Biosciences) supplemented with 10% OADC (BD Biosciences), 0.5% glycerol (Fisher), and 0.1% Tween-80 (Fisher) or on 7H10 plates. All work with M. tuberculosis was performed under Biosafety level 3 containment using procedures approved by the Texas A&M University Institutional Biosafety Committee. For Mtb infections, the inoculum was prepared by growing bacteria to log phase (OD600 0.6-0.8).

Bacterial cultures were spun at low speed (500 rpm) for 5 minutes to remove clumps. Bacteria were then pelleted with a spin at 3000 rpm 5 min and washed with 1X PBS three times. Upon final wash, the resuspended bacteria were briefly sonicated and spun at low speed once again to further remove clumps. The bacteria were diluted in DMEM (Hyclone) + 10% horse serum (Gibco) in vitro infections, or 1X PBS in vivo infections. For *in vitro* infections, plates containing bacteria and cells were spun for 10minutes at 1000 rpm to synchronize infection. The cells were washed with 1X PBS two additional times before the cells were given fresh media.

### *In vivo* Bacterial Infections

The Mtb inoculum was prepared as described above. Age-and sex-matched mice were infected via inhalation exposure using a Madison chamber (Glas-Col) calibrated to introduce 100-200 CFUs per mouse. For each infection, approximately 5 mice were euthanized immediately, and their lungs were homogenized and plated to verify an accurate inoculum. Infected mice were housed under BSL3 containment and monitored daily by lab members and veterinary staff. At the indicated time points, mice were euthanized, and tissue samples were collected. Organs were divided to maximize infection readouts (CFUs: left lobe lung and ½ spleen; histology: superior right lobe and 1/2 spleen). For histological analysis organs were fixed for 24 hours in neutral buffered formalin and moved to 1X PBS (lung, spleen). Organs were further processed as described below.

### Histopathology

Mtb-infected mouse lungs were fixed with 10% neutral formalin and processed, embedded in paraffin, and cut into 5 µm sections and stained with H&E or AFB (AML Laboratories). A boarded veterinary anatomic pathologist performed a blinded evaluation of lung sections for inflammation. To quantify the percentage of lung fields occupied by histiocytic inflammatory infiltrates, scanned images of a lobe of each lung were analyzed using QuPath Bioimage analysis version 0.4.3 to determine the total cross-sectional area of inflammatory foci per total lung cross-sectional area. Additional criteria were evaluated by dividing the digital images into 500 × 500 µm grids and counting the percentage of squares containing neutrophils, neutrophils arranged in clusters (>5 neutrophils in close approximation), necrotic debris, and foamy macrophages.

### Immunoprecipitation and Mass spectrometry

1.8 × 106 HEK293T cells in a 10cm plate were transfected with 1-5 µg of 2xSTREP-Rv1075c or 2xSTREP-GFP using PolyJet In Vitro DNA Transfection Reagent. 24 hours post transfection, cells were harvested, washed with PBS, and collected using PBS containing 10 mM EDTA. Cell pellets were lysed in ice-cold lysis buffer (50 mM Tris-HCl, pH 7.4, 150 mM NaCl, 1 mM EDTA, 0.075% NP-40) supplemented with EDTA-free protease and phosphatase inhibitors (thermo 78447). Following centrifugation (14,000 rpm, 10 min, 4°C), the nuclear pellet was resuspended in lysis buffer supplemented with EDTA-free protease and phosphatase inhibitors, sonicated (30 s on/30 s off cycles for 30–40 min), and clarified by centrifugation. An aliquot of the nuclear lysate was retained as the input control. Nuclear lysates were incubated with streptavidin beads for 60 min at 4°C with end-over-end rotation. Beads were washed three to five times with wash buffer (50 mM Tris-HCl, pH 7.4, 150 mM NaCl, 1 mM EDTA, 0.05% NP-40), and bound proteins were competitively eluted with biotin (10mM biotin resuspended in IP lysis buffer) three times at 20 minute intervals at 4°C. Eluates were combined. For mass spectrometry, protein lysates were submitted to the UT Southwestern Proteomics Core Facility for sample preparation and LC-MS/MS analysis. Samples underwent disulfide bond reduction, alkylation, trypsin digestion, and peptide cleanup prior to analysis on a Q Exactive HF mass spectrometer. Protein identification and quantification were performed by the UT Southwestern Proteomics Core Facility using their standard workflows. For western blotting, the protein lysates were mixed with SDS sample buffer containing β-mercaptoethanol, boiled for 5 min, and analyzed by SDS-PAGE and immunoblotting.

### Plate-based assays for cell death

BMDMs were plated in 96-well clear bottom plates (Corning) at 2.5 × 10^4^ cells/well in 50 µL of media. Following cell adherence (1 h), an additional 25 µL was added to each well. The following day, media were removed, and respective agonists were used to initiate various cell death modalities. For cell death assays, 5 µg/mL PI (ThermoFisher) was added at the time of adding cell death agonists. Total cell numbers used for normalization were counted on a subset of cells using NucBlue (ThermoFisher) in 1× PBS. Live cell imaging was done using 4× magnification on a Lionheart plate reader. Post-run analysis was conducted using Gen5 version 3.15 (Biotek) software.

### Immunofluorescence

BMDMs were seeded at 3×10^5^ cells/well on glass coverslips in 24-well dishes. Raw 264.7 macrophages were seeded at 2.5×10^5^ on glass coverslips treated for 20 minutes with poly-l-lysine (Fisher 19321-A) in 24-well dishes. Cells were fixed in 4% PFA for 15 min at RT and then washed three times with PBS. Coverslips were incubated in primary antibody diluted in PBS + 5% non-fat milk + 0.1% Triton-X (PBS-MT) for 3 hours in the dark. Primary antibodies used in this study were Ms-strep (Genscript, cat A01732-100), Rb-strep (Genscript A00626-40), Rb-Wdr5 (cell signaling13105s), Rb-Rbbp5 (cell signaling 13171s), Rb-Ash2l (abcam ab176334), Rb-Dpy30 (bethyl a304-296a), Ms-Mtb (Novus Bio NB100-62769), or Rb-Nucleolin (Abcam ab22758). Cells were then washed three times in PBS and incubated in secondary antibodies (invitrogen goat anti-rabbit Alexa Fluor 488, goat anti-mouse Alexa Fluor 488, or goat anti-mouse Alexa Fluor 594 diluted 1:500 in PBS for 1 hr in the dark. When stated, lipidspot (Biotium, 70065-T) was incubated on coverslips for 10 minutes in the dark. Coverslips were washed twice with 1X PBS and twice with deionized water and mounted on glass slides using Fluoromount-GTM with DAPI (Invitrogen 00-4959-52). Z-stack images were obtained using an Olympus IX83 inverted confocal microscope or a Zeiss LSM 710 confocal microscope.

### Protein immunoblot

Cells were washed with PBS and lysed in 1X RIPA buffer with protease and phosphatase inhibitors, with the addition of 1 U/mL Benzonase to degrade genomic DNA. Proteins were separated by SDS-PAGE and transferred to nitrocellulose membranes. Membranes were blocked for 1 hr at RT in LiCOR Odyssey blocking buffer or TBS with 5% BSA. Blots were incubated overnight at RT with the following antibodies: were Ms-ACTB (Abcam, 1:2000) Ms-strep (Genscript, cat A01732-100), Rb-strep (Genscript A00626-40), Rb-Wdr5 (cell signaling13105s), Rb-Rbbp5 (cell signaling 13171s), Rb-Ash2l (abcam ab176334), Rb-Dpy30 (bethyl a304-296a). Membranes were incubated with appropriate secondary antibodies for 2 hrs at RT prior to imaging on a LiCOR Odyssey Fc Dual-Mode Imaging System.

### Quantification and statistical analysis

All data are representative of two or more independent experiments with n=3 or greater unless specifically noted in the figure legends. For all quantifications, n represents the number of biological replicates, either number of wells containing cells, number of mice, or number of flies. Error bars represent STDEV. For *in vitro* assays, statistical significance was determined using either a two-tailed Student’s unpaired T test, one-way ANOVA with Sidak’s post hoc test, or two-way ANOVA with Tukey’s post hoc test. For *in vivo* mouse infections, significance was determined using a Mann-Whitney U test based on the assumption that samples (mice) followed a non-normal distribution. The specific statistical test used to determine significance for each experiment is listed at the end of the figure legends.

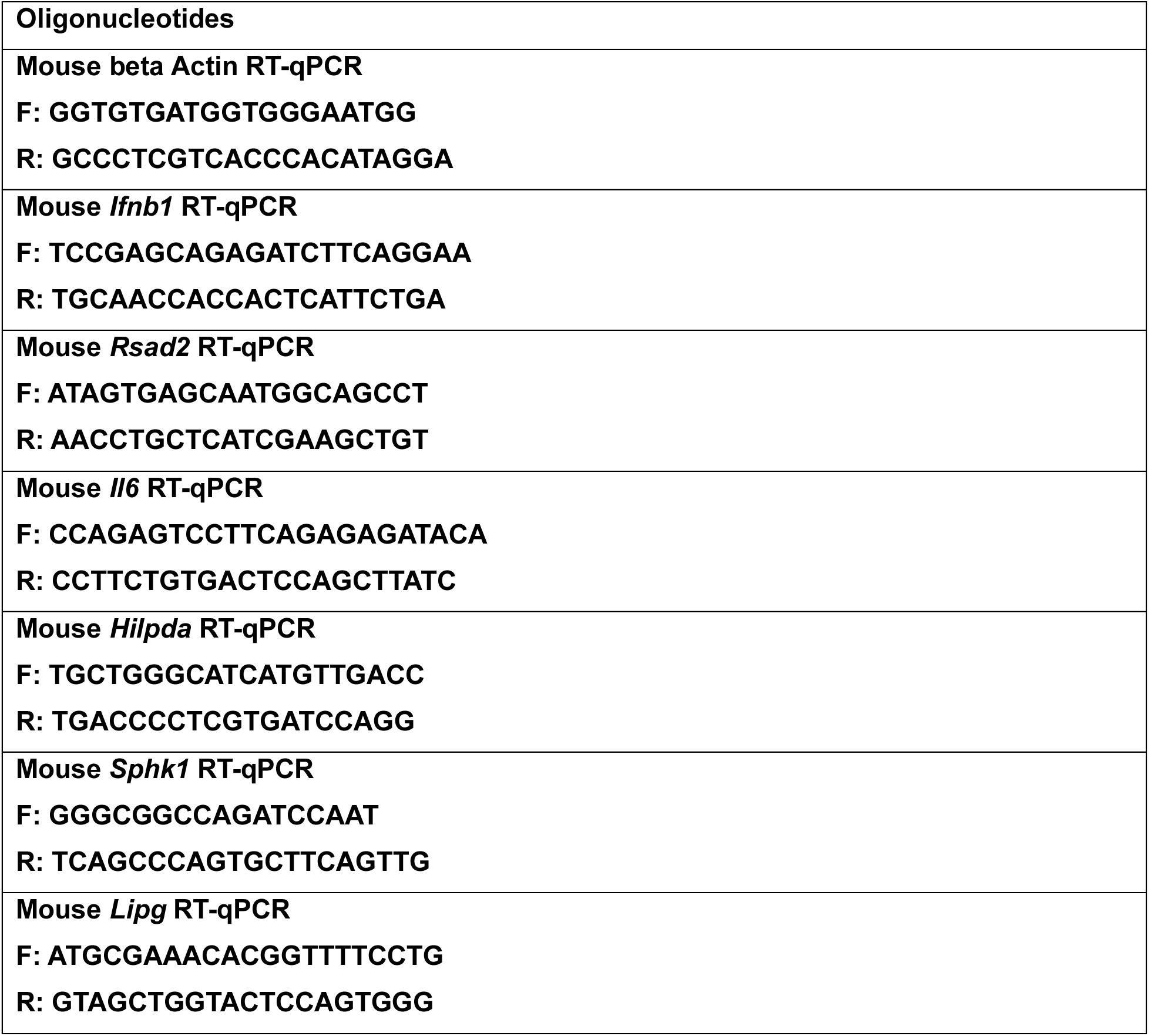
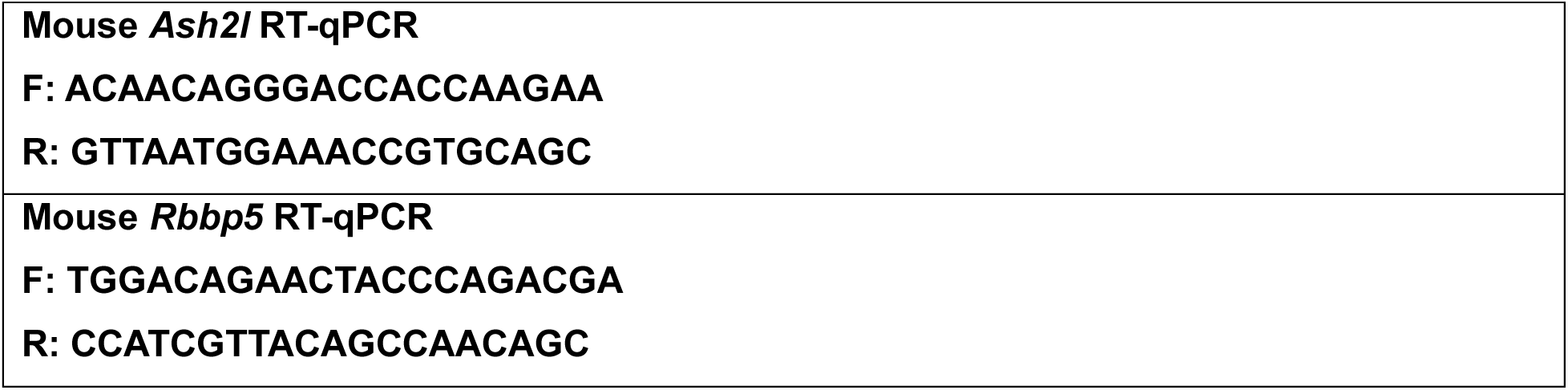

