## Supplemental figures and legends for "*Mycobacterium tuberculosis* manipulates host inflammation and lipid metabolism through the SET1-interacting protein Rv1075c"

A

| Mtb | Score | Amino Acid Residues | Motif |
|---|---|---|---|
| Rv2823c | 3.8 | GTDRRKADSDDGHGASTWDPDTPLYSMFNR | Classical Bipartite |
| Rv3899c | 3.8 | VRKTGVLENEAELLHGCITAVKESVLKAYP | Classical Bipartite |
| Mce1d | 3.7 | DKVQIMGLPVGSIDKIEPAGDKMKVTFHY | Classical Bipartite |
| Caea | 3.6 | TLPKRVHERFDLVGFDPRGVASSRPAIWCNS | Classical Bipartite |
| LpqR | 3.6 | RQCPSVRRSGELCLADMGTDFDDFSSRATAFAT | Classical Bipartite |
| Rv2075c | 3.5 | RPNPARRATNGCVPLPLDVSREEIRASGARAV | Classical Bipartite |
| EspB | 3.5 | SKGSQQEDEALYTEDRAWTEAVIGNRRR | Classical Bipartite |
| Rv3722c | 3.5 | RRTVALAKDVGIAVTEAGASFPYRKDPDDKNIRI | Classical Bipartite |
| Rv2469c | 3.4 | GKKRRGHRSSGVAAGVTGPASCLHSVHSH | Classical Bipartite |
| PPE11 | 3.4 | RRRRRAAAKERGNADEFVDMDSGPAIPPSGERDAW | Classical Bipartite |
| Apa | 3.3 | TRRKGRLAALAIAAMASASLVTVAVPATAN | Classical Bipartite |
| Rv3369 | 3.3 | RDDAPYWAKYREDAAKFGLTEAIAAYSTRLKITPT | Classical Bipartite |
| Rv0787 | 3.3 | RPPWLAQLRRRLRIGVQLGSRVVLEQGRQPRDVYVI | Classical Bipartite |
| Rv3267 | 3.2 | RIRKRLSRGVMTLVSVVALLMTGAGYWVAH | Classical Bipartite |
| Mce1b | 3.1 | RYVELKRGEGKGANDLLPPGGLIPLSRTSPALD | Classical Bipartite |
| LpqB | 3.1 | ETYKRNTLYFADPTGKTVVPDPRYVAVSDRD | Classical Bipartite |
| EspR | 3.1 | FFRIKAAYFTDDEYYEKLDKELQWLCTMRDD | Classical Bipartite |
| Rv3491 | 3.1 | QPKLSKQPFSLQLIGPPPSPVQRYPLYCN | Classical Bipartite |
| Tb18.6 | 3 | KVEKLDLPEDASPAYLGFNLFQHAIARAVI | Classical Bipartite |
| FbpA | 3 | GKAGCQTYKWETFLTSELPGWLQANRHVK | Classical Bipartite |
| Rv3627c | 3 | RRSRTPALDAGRELAKALGLDPAAVTIASAP | Classical Bipartite |

B

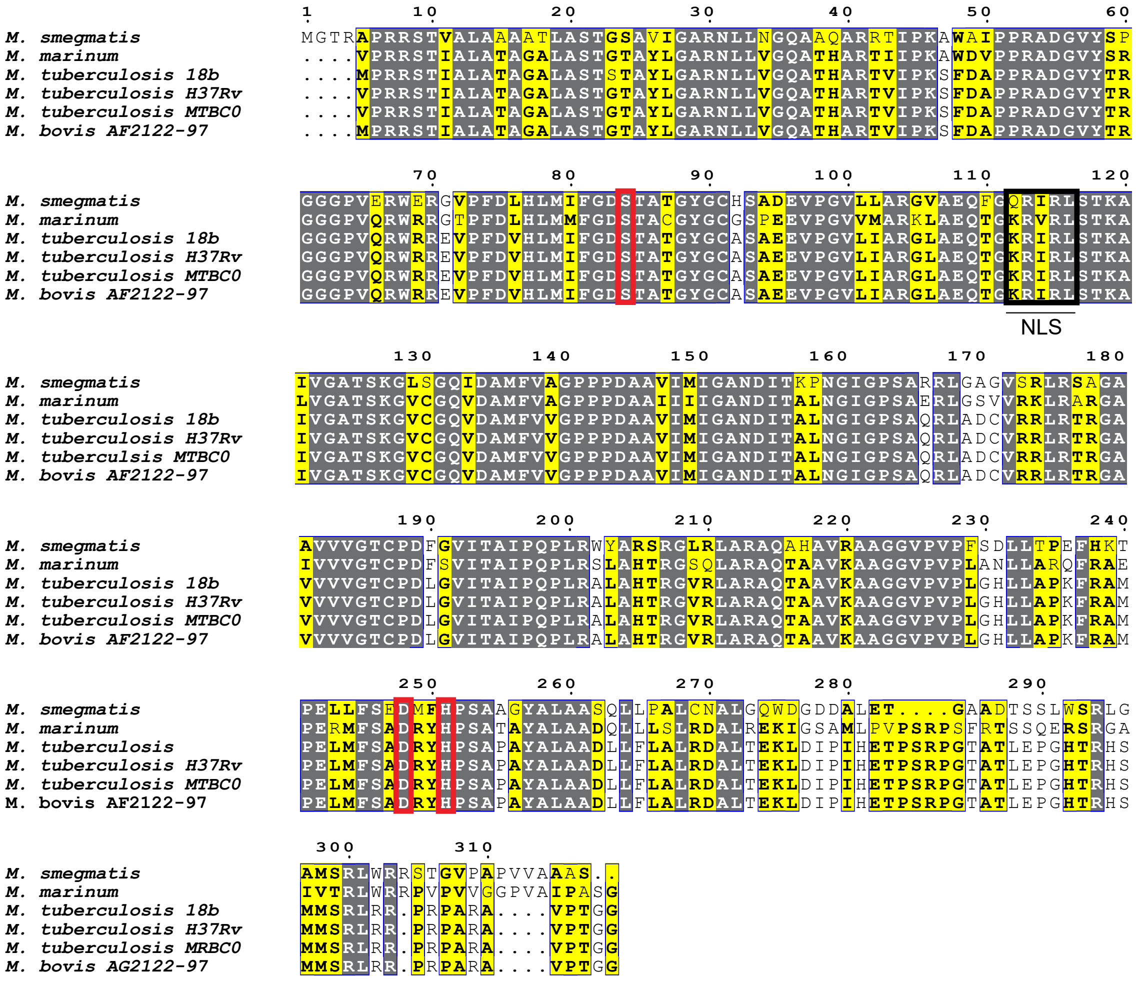


Figure S1

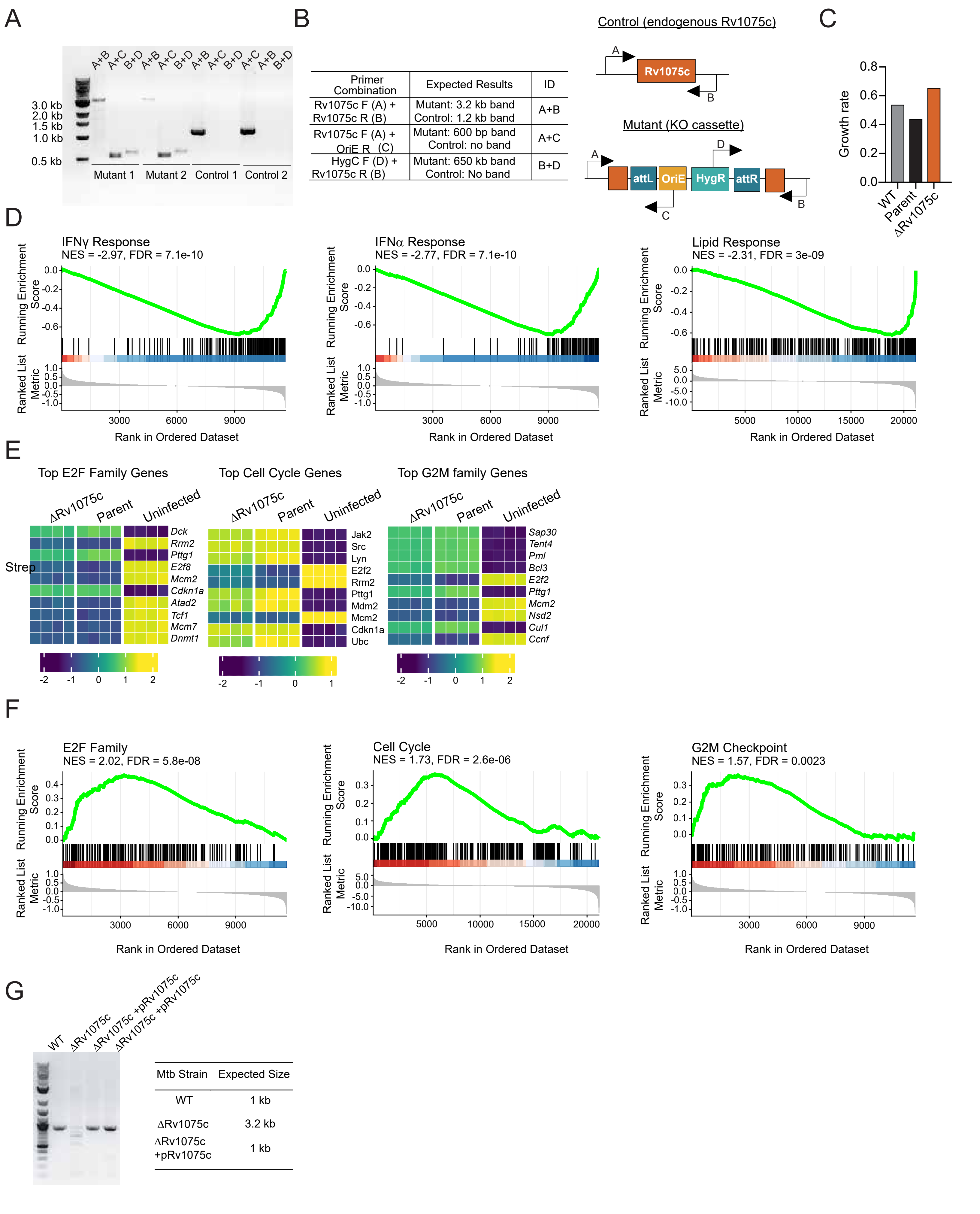
A
A+B A+C B+D A+B A+C B+D A+B A+C B+D A+C A+B B+D
3.0 kb
2.0 kb
1.5 kb
1.0 kb
0.5 kb
Mutant 1
Mutant 2
Control 1
Control 2
B
Primer Combination | Expected Results | ID
Rv1075c F (A) + Rv1075c R (B) | Mutant: 3.2 kb band Control: 1.2 kb band | A+B
Rv1075c F (A) + OriE R (C) | Mutant: 600 bp band Control: no band | A+C
HygC F (D) + Rv1075c R (B) | Mutant: 650 kb band Control: No band | B+D
Control (endogenous Rv1075c)
Rv1075c
Mutant (KO cassette)
attL
OriE
HygR
attR
C
Growth rate
WT
Parent
ΔRv1075c
D
IFNγ Response
NES = -2.97, FDR = 7.1e-10
IFNα Response
NES = -2.77, FDR = 7.1e-10
Lipid Response
NES = -2.31, FDR = 3e-09
Running Enrichment Score
Ranked List Metric
Rank in Ordered Dataset
E
Top E2F Family Genes
Top Cell Cycle Genes
Top G2M family Genes
ΔRv1075c
Parent
Uninfected
Strep
Dck
Rrm2
Pttg1
E2f8
Mcm2
Cdkn1a
Atad2
Tcf1
Mcm7
Dnmt1
Jak2
Src
Lyn
E2f2
Rrm2
Pttg1
Mdm2
Mcm2
Cdkn1a
Ubc
Sap30
Tent4
Pml
Bcl3
E2f2
Pttg1
Mcm2
Nsd3
Cul1
Ccnf
F
E2F Family
NES = 2.02, FDR = 5.8e-08
Cell Cycle
NES = 1.73, FDR = 2.6e-06
G2M Checkpoint
NES = 1.57, FDR = 0.0023
Running Enrichment Score
Ranked List Metric
Rank in Ordered Dataset
G
WT
ΔRv1075c
ΔRv1075c +pRv1075c
ΔRv1075c +pRv1075c
Mtb Strain | Expected Size
WT | 1 kb
ΔRv1075c | 3.2 kb
ΔRv1075c +pRv1075c | 1 kb

Figure S2

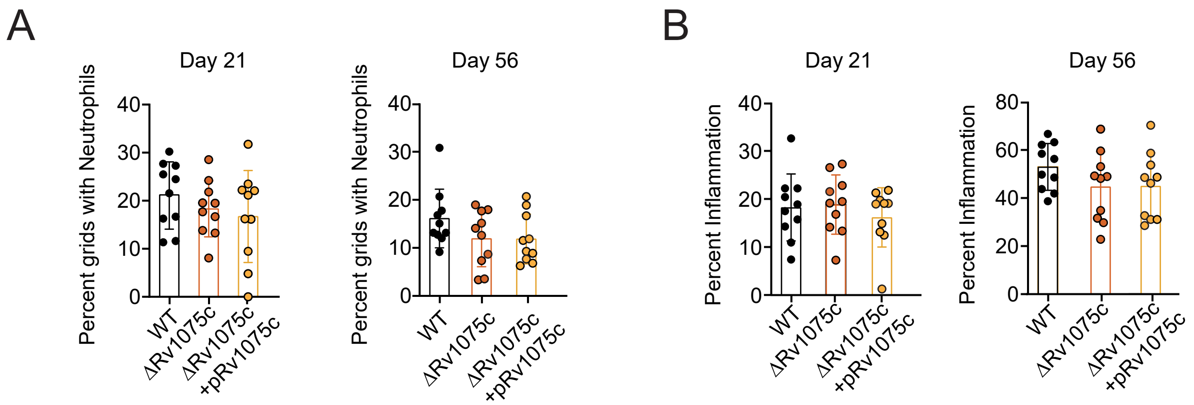
A
Day 21
Day 56
Percent grids with Neutrophils
0
10
20
30
40
WT
ΔRv1075c
ΔRv1075c +pRv1075c
B
Day 21
Day 56
Percent Inflammation
0
10
20
30
40
60
80

Figure S3

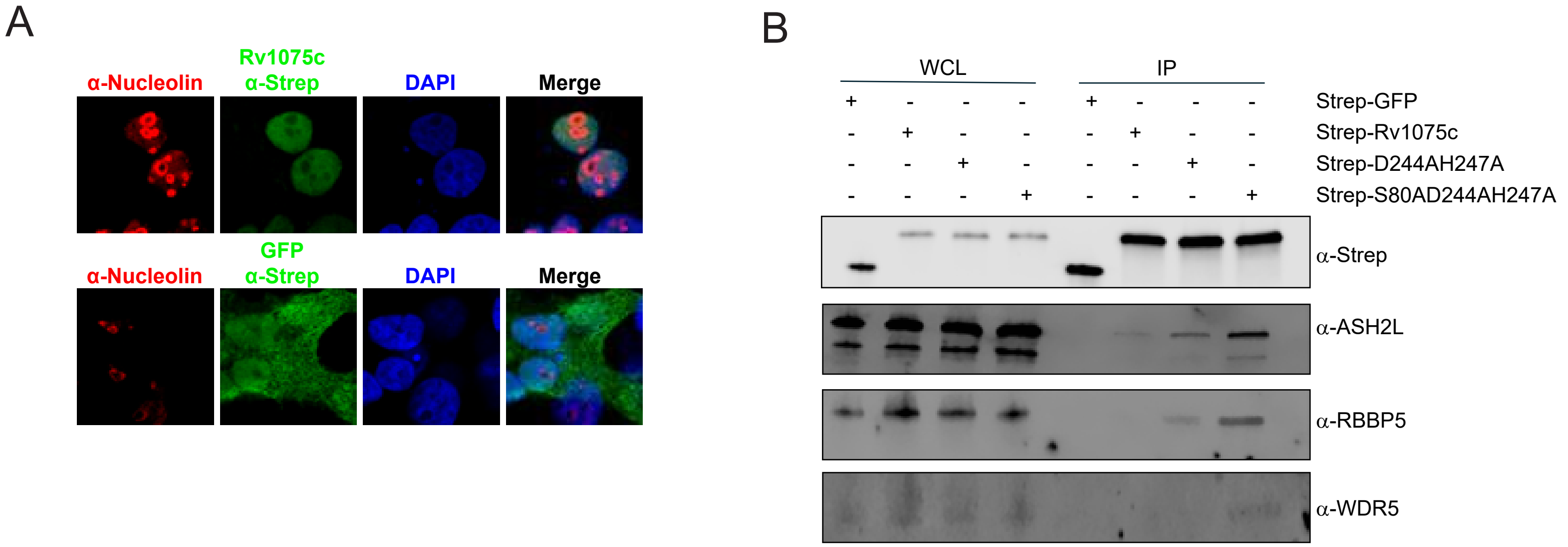
A
α-Nucleolin
Rv1075c
α-Strep
DAPI
Merge
α-Nucleolin
GFP
α-Strep
DAPI
Merge
B
WCL
IP
Strep-GFP
Strep-Rv1075c
Strep-D244AH247A
Strep-S80AD244AH247A
α-Strep
α-ASH2L
α-RBBP5
α-WDR5

Figure S4

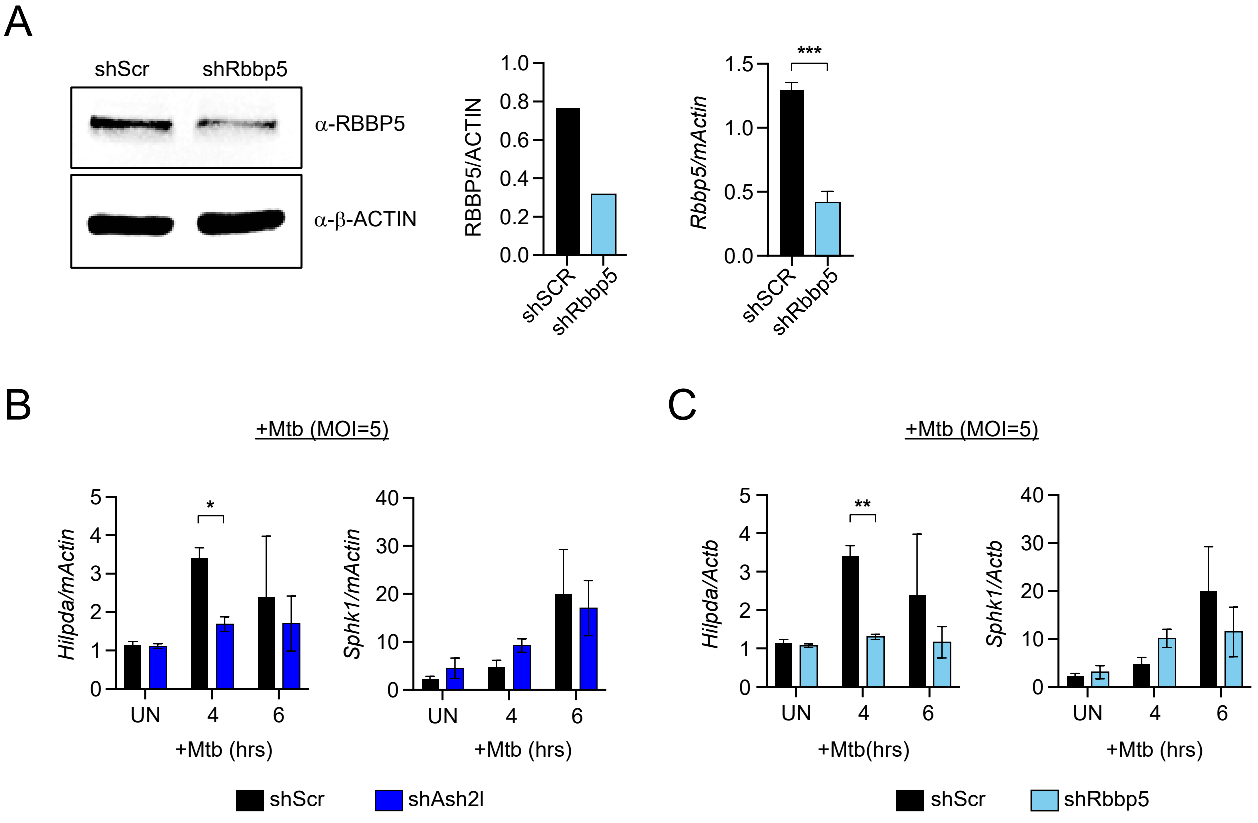
A
shScr
shRbbp5
α-RBBP5
α-β-ACTIN
RBBP5/ACTIN
1.0
0.8
0.6
0.4
0.2
0.0
shSCR
shRbbp5
Rbbp5/mActin
***
1.5
1.0
0.5
0.0
shSCR
shRbbp5
B
+Mtb (MOI=5)
Hilpda/mActin
*
Sphk1/mActin
UN
4
6
+Mtb (hrs)
shScr
shAsh2l
C
+Mtb (MOI=5)
Hilpda/Actb
**
Sphk1/Actb
UN
4
6
+Mtb(hrs)
+Mtb (hrs)
shScr
shRbbp5

Figure S5

**SUPPLEMENTAL FIGURE LEGENDS**

**Figure S1:**

A. Alignment of Rv1075c orthologs from *M. tuberculosis* H37Rv (Rv1075c), *M. tuberculosis* 18b (MT18B_1419), *M. tuberculosis* MTBC$_0$ (mtbc0_001155), *M. smegmatis* MSMEG_5272*, M. bovis* AF2122/97 (Mb1104c), and *M. Marinum (MMAR_4392).* Predicted NLS is underlined.

**Figure S2:**

A. Agarose gel electrophoresis (2.5%) of PCR amplified products using primers specific for the verification of ORBIT mutants. Lanes 1 represents ladder, lanes 2-4 represent mutant 1, lanes 5-7 represent mutant 2, lanes 8-10 represent control 1, lanes 11-13 represent control 2.

B. Table and diagram depicting primer design strategy to amplify Rv1075c and/or segments of payload plasmid.

C. Bar graph representing the doubling rate of WT (grey), the ORBIT Parental strain (black), and the ΔRv1075c (orange) mutant in liquid culture.

D. Gene set enrichment analysis (GSEA) for IFNg response, IFNa response, and Lipid response of ΔRv1075c vs Parental WT infected macrophages from RNA-seq experiments using differentially expressed genes (DEG). Normalized enrichment scores (NES) and FDR values are shown for each GSEA.

E. Heatmap derived from ΔRv1075c vs Parental WT infected BMDMs RNA-seq of top enriched

genes from the pathways the top E2F family pathway, cycle cell genes, and G2M family genes.

F. Gene set enrichment analysis (GSEA) of E2F family, Cell cycle, and G2M checkpoint of ΔRv1075c vs Parental WT infected macrophages from RNA-seq experiments using differentially expressed genes (DEG). Normalized enrichment scores (NES) and FDR values are shown for each GSEA.

G. Agarose gel electrophoresis (2.5%) of PCR amplified products using primers specific for the verification of the complementation of Rv1075c in ORBIT dRv1075c mutant. Lane 1 represents ladder, lane 2 represents ORBIT mutant ΔRv1075c, lanes 3-4 represent complemented ΔRv1075c +pRv1075c.

**Figure S3:**

A. Percent neutrophil-containing grids infiltration in Mtb infected lungs of WT, DRv1075c, and DRv1075c+pRv1075c infected mice at Day 21 and 56 post infection.

B. As in A but for % inflammation

**Figure S4:**

A. IF microscopy of Rv1075c (green, anti-strep), nucleolin (red, anti-nucleolin), and DAPI (blue) in HEK293T cells ectopically expressing C-2xSTREP-Rv1075c or C-2xSTREP-GFP control.

B. Immunoblot of Strep-IP of Rv1075c, GFP, $D^{244A}$-$H^{247A}$, or $S^{80A}$-$D^{244A}$-$H^{247A}$ probed for endogenous Ash2l (anti-Ash2l), Wdr5 (anti-Wdr5), and Rbbp5 (anti-Rbbp5) from HEK 293T cells

**Figure S5:**

A. Protein and transcript expression of Rbbp5 (anti-Ash2l) in shScr and shRbbp5 RAW 264.7 macrophages. Student T test. Error bars represent STDEV. **$p<0.005$

B. RT-qPCR of *Hilpda* and *Sphk1* in shAsh2l (dark blue) and shScr (black) RAW 264.7 macrophages infected with Mtb for 4 and 6 hours. Two-way Anova. Error bars represent STDEV. *$p<0.05$.

C. As in B but in shRbbp5 (light blue) and shScr (black) RAW 264.7 macrophages
